# Lymphatics regulation of the inflammatory clotting creates the natural on-off switch for the immune ignorance that allows subcutaneous allografting

**DOI:** 10.1101/2021.06.22.449446

**Authors:** Małgorzata Wachowska, Witold W Kilarski

## Abstract

The ability of lymph to clot indicates that, like blood vessels, lymphatics must have means to counteract this process. Here, we analyzed lymphatic hemostatic properties, tailoring them for potential therapeutic applications. Inflammatory stimuli induced tissue factor-dependent focal lymph clotting while blocking thrombomodulin leading to widespread but transient occlusion of collecting vessels. Decellularization of lymphatics resulted in tissue factor-independent lymphatic occlusion by widespread and persistent lymph clots. In occluded decellularized ‘ghost’ vessels, fibrin was eventually reperfused. During the regeneration, ghost lymphatics were filled with granuloma-like clusters of antigen-presenting cells and T cells. Despite that, immune response against allografts placed under non-drained skin did not develop as long lymphatics remained occluded, the effect that could be prolonged by delaying regeneration of the decellularized collectors. When the lymph clotting was blocked, decellularized lymphatics could still drain macromolecules and leukocytes, showing that lymphatic endothelium is not necessary for the classic lymphatic functions. The control of excessive clotting emerges as the essential function of lymphatics that could explain the seeming spandrel presence of lymphatic networks in organs such as the kidney or heart, contribute to microvascular thrombosis during infection, and can be exploited to induce immune ignorance of the subcutaneous endocrine grafts.

## Introduction

The Intima lining of blood vasculature forms the only known surface that dynamically controls blood liquidity(Gimbrone,1987). Despite ontogenetic and functional similarities, that function has not been yet ascribed to its relative circulation of lymphatic vessels. However, since the first description of the lymphatic system by Olaf Rudbeck over 350 years ago, lymph is known for its ability to clot outside of the lymphatic vessels (discussed in (Rusznyák,& al,1967), (Drinker,& al,1933), (Miller,& al,2000)). However, even though lymph clotting time is slower as compared to blood, it has intrinsic clotting properties(Opie,1913, Copley,& al,1942)(Fantl,& al,1953). Lymph clotting time varies in a broad, up to 20-times range as a concentration of lymph clotting factors that fluctuate between species, individuals, physiological conditions or even sampling location, and is 10 to 70% lower as compared to the blood (Mayanskii,& al,1966, Le,& al,1998, Leak,& al,2004). Together with the absence of platelet-derived anionic phospholipids required for activation of factor X and prothrombinase and a higher concentration of fibrinolytic factors, lymphatics assure lymph hypocoagulability under physiological conditions (Fantl,& al,1953, Lippi,& al,2012). Also, due to the lack of thrombocytes, lymph, in contrast to blood clots, did not retracted(Fantl,& al,1953), producing transparent clots that are similar to the arterial thrombus. On the other hand, relatively slow lymph flow and a possibility to substitute platelet phospholipids with membrane components of endothelial cells (Brinkman,& al,1996) may favor lymph clotting during inflammation, when augmented blood vessel permeability to plasma proteins leads to 3-fold enrichment of interstitial fluid and lymph(Drinker,& al,1933) when protein extravasation from blood increases up to 100 times during pathological conditions(Rusznyák,& al,1967, Aukland,& al,1993, Wiig,& al,2012).

These observations confirm that lymph is a hemostatic fluid. More importantly, it infers that like blood vessels, lymphatics must have specific anti-coagulatory properties that enable them to control lymph fluidity. By doing so, the lymphatic system could regulate other passive functions of lymphatics, i.e., fluid drainage and immune cell trafficking, and for instance, limit the systemic spread of parasites and infections(Engelmann,& al,2013). However, only a handful of studies address thrombo-regulatory properties of lymphatic endothelium (for review see (Lippi,& al,2012)). For example, it was shown that similar to blood endothelial cells, lymphatic collectors express thrombomodulin, a membrane receptor capable of switching the substrate specificity of thrombin to protein C, thus making thrombin a potent anticlotting factor(Conway,2012). Importantly, the lymphatic endothelium does not produce the von Willebrand factor, a multimeric protein that participates in platelet activation on the exposed sub-endothelial collagen. This phenotype is in line with the lack of platelets in lymph(Maruyama,& al,1985). Such a specificity in a loss of the non-essential factor of pro-thrombotic system suggesting that lymphatic anti-hemostatic properties were preserved not just as an evolutionary spandrel(Gould,& al,1979), a leftover heritage from ontogenetically ancestral blood endothelial linage(Jafree,& al,2021).

In response to pro-inflammatory molecules such as cytokines TNFα, IL-1β, or TLR agonist LPS, blood endothelium can turn into a pro-thrombotic surface, e.g., by initiating the expression of tissue factor (TF), membrane receptor initiating extrinsic coagulation pathway(Kirchhofer,& al,1994). Few *in vitro* studies have demonstrated that lymphatic endothelial cells are also capable of responding to inflammatory stimuli, for example, with increased synthesis of plasminogen activators and inhibitors (Lippi,& al,2012, Laschinger,& al,1990). In contrast to blood circulation, where vessel occlusion by thrombus leads to tissue ischemia and immediate localized symptoms, such as pain and necrosis, lymphedema following lymphatic blockage can take years to develop and requires additional factors to precipitate the condition(Drinker,& al,1933). Yet, there is clinical evidence of pathological lymph clotting that accompanies specific inflammatory pathologies. For example, intralymphatic lymph clots encapsulating the granuloma were found in the chronic inflammation(Olszewski,2002), surrounding dead nematodes in lymphatic filariasis(Case,& al,1991, Fader,& al,1986), and occluding vessels in patients with idiopathic primary lymphedema(Hara,& al,2013) or secondary lymphedema (Cluzan,2007). These clots can undergo fibrosis, forming long-lasting sclerotic intravascular intrusions in chronic diseases(Olszewski,2002, Fader,& al,1986, Cluzan,2007). The tumor microenvironment is another typical inflammatory pathology, where leaky blood vessels and chronic inflammation should constitute ideal conditions for intravascular lymph clotting. Indeed, biomarkers of tumor stroma, i.e., TF expression, interstitial fluid hypercoagulability, and stromal fibrin deposition(Forster,& al,2006, Ruf,& al,2011) correlate with dysfunctionality or complete collapse of intra-tumoral lymphatics in most human cancers(Padera,& al,2002). However, because fibrin, as a provisional matrix rapidly undergoes remodeling and digestion (Idell,2003) while pre-symptomatic tumor development is a chronic disease that can take years to develop. Therefore, it is not surprising that there are only scant reports describing fibrotic occlusion of lymphatics associated with human tumors (Margaritescu,2015) or as identified by us in an experimental mouse model of B16-F10 melanoma (Supplementary Fig. 1).

The transport of clottable lymph requires anchoring of anti-clotting mechanism in the walls of lymphatic collectors but also creates an opportunity for lymphatic endothelium to dynamically change vessel patency depending on tissue condition. Therefore, it should be possible to reverse the anti-hemostatic properties of lymphatics with pharmacological inhibition of thrombomodulin, turning the lymphatic endothelium into a pro-thrombotic surface with, e.g., the expression of pro-thrombotic tissue factors or decellularization of lymphatic collectors. The latter would remove the endothelium-derived anti-clotting factors, such as thrombomodulin and plasminogen activators but also exposes negatively charged basement membranes(van der Meijden,& al,2009) that could activate intrinsic coagulation pathways(White-Adams,& al,2010). Surprisingly, we found that intact lymphatic endothelium was indispensable for lymph and immune cell transport through collecting vessels. As proof that the anti-clotting function of lymphatic endothelium can be applied therapeutically, we showed that a complete block of lymph transport to the draining lymph node blocked the immune response against subcutaneous allograft until drainage of lymphatics was restored. By blinding the lymph node to graft antigens with a local treatment we could prolong the survival of subcutaneously allografted tissues without systemic and toxic immunosuppression.

Remarkably, the lack of anti-coagulatory lymphatic drainage might contribute to the delayed organ failure as lymphatic circulation is never reinstated during kidney or heart transplantation(Shoskes,2011). Similarly, the abnormalities in the anti-thrombotic system of lymphatics could explain the mysterious pathology in COVID-19 patients, when the decrease in viral load correlates with the increase in blood micro clots leading to failure of organs that were never infected(Fahmy,& al,2021).

## Results

### Induced expression of TF leads to lymph clotting and obstruction of drainage

Here we analyzed the anti-clotting properties of lymphatics. In a physiological state, the endothelium of the lymphatic collector, like blood endothelial cells, expressed an anti-thrombotic thrombomodulin receptor and did not express pro-coagulatory tissue factor (TF, **Fig. 1A**). Thrombomodulin expression on lymphatics could not be just a spandrel(Gould,& al,1979), a left-over expression pattern heritage from the ancestral blood endothelial lineage. This is because contrary unlike blood, lymphatic vessels, which are not expected to encounter platelets, also lack the Weibel-Palade stores of von Willebrand factor, the ligand that activates platelets in the vascular wounds (**Fig. 1A, right**). Further, we tested whether the anti-coagulatory environment of the lymphatic could be reverted, and the stimulation of *de novo* expression of TF on lymphatic endothelium appeared as the most intuitive approach. To assay the activation of intralymphatic coagulation, we intravenously injected fluorescently labeled fibrinogen (**Fig. 1B**). If fibrinogen leaked to the tissue and was polymerized into insoluble fibrin, its fluorescent deposits could be detected within thin mouse ear dermis with intravital microscopy or, after fixation, in the whole ear skin preparation with confocal optical sectioning.

**Figure 1.**
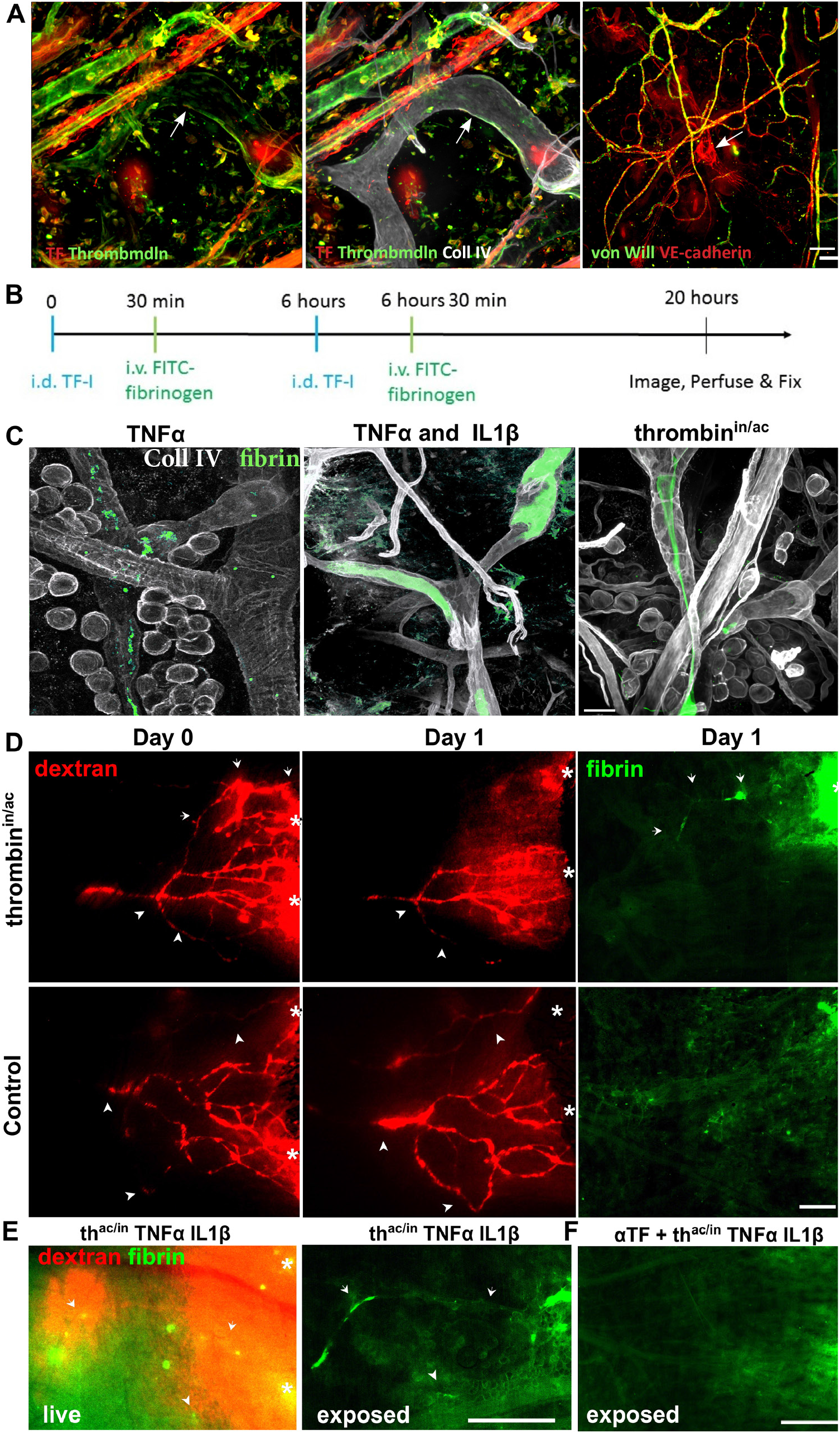
Inflammatory mediators lead to lymph coagulation at selected collector. **A. Left**: The pattern of thrombomodulin expression overlapped with the expression of endothelial marker CD31 but was also present on some perivascular and interstitial cells but also within nerve fibers (n). Arrows point to collecting lymphatics that have unique morphological features, i.e., the presence of valves with uneven vessel diameters. **Middle**: In non-inflamed ear skin, endothelial cells of lymphatic collectors similar to blood endothelium express thrombomodulin but not tissue factor (TF), whose expression is restricted to perivascular cells and adipocytes (a). **Right:** In contrast to blood endothelium, lymphatic cells were negative for the von Willebrand factor. Arrows point to lymphatic collectors. **B**. Schematic of experimental design. Intradermal injection of known stimulators of TF expression (TNFα, IL1β and thrombin^in/ac^ (partially inactivated thrombin) on day 0 was followed by intravenous injection of FITC-labeled fibrinogen. Six hours after the first injection, the procedure is repeated. The mouse is perfused and fixed 13 hours later. **C**. Two subsequent remote intradermal injections of TF-I TNFα, TNFα, and IL1β or partially inactivated thrombin (thrombin^in/ac^, a mix of proteolytically inactive and active thrombin; 6:1 ratio) resulted in the formation of intralymphatic clots within specific locations of lymphatic collectors as shown by the presence of fluorescent fibrin. Basement membrane was stained by collagen IV. **D, Top:** Demonstration of lymphatic drainage by intradermal injection of fluorescent dextran. Two consecutive injections (6 hours apart on day 1) of thrombin^in/ ac^ blocked lymphatic drainage in some branches of the collecting lymphatic network 12 hours after the second injection. Intralymphatic clots (fibrin) are present in with the location of lymphatic collector junction functional lymphatic was blocked (arrows). **D, Bottom:** Intradermal injections of PBS had no adverse effect on lymphatic perfusion. Arrows point to collectors that were blocked by intralymphatic fibrin (green). Arrowheads mark the same functional vessels visualized on the day of the experiment (Day 0) and the following day. **E, Left**: Superficial view of the live dorsal skin of the ear after lymphangiography. Combination of all three pro-clotting factors in single treatment had no cumulative effect and, similar to the individual-factor treatment, induced lymph clotting that blocked vessel drainage only in certain collectors. th^at/in^ – partially inactivated thrombin. **E, Middle**: Internal view of the same ear skin after the mouse was fix-perfused and dorsal skin exposed by detaching ventral skin flap with cartilage. Arrowhead and arrow point to the same vessel locations. **E, Right**: Internal view of the exposed dorsal dermis where lymph clotting was prevented with co-injection of tissue factor-blocking antibody (αTF). The intensity of the von Willebrand factor was normalized to its signal level in capillaries. The intensity of lymphatic fibrin deposits was normalized to the signal level of clots at the sites of injection (*). Scale bar: **A, C** – 50μm, **D, E** – 250μm.collectors. Arrows in D point to lymphatic occluded by fibrin clots. Stars (*) in D and E mark sites of intradermal injection of TRITC-dextran. Scale bar: 50μm.

TNFα and IL1β are cytokines known as potent inducers of tissue factor (TF) expression *in vitro* (Kirchhofer,& al,1994, Bevilacqua,& al,1984) but only on subsets of blood endothelial cells *in vivo* (Lupu,& al,2005). Inflammatory stimulation of TF on the intima layer of blood vessels results in activation of the extrinsic pathway within vessel lumen and temporary, local vessel occlusion by thrombus. Similarly, in our study, even in the absence of platelets, repeated intradermal injections of TNFα induced clot formation within draining collectors but only at discrete locations **(Fig.1C)**. No cumulative effect was observed when TNFα was co-injected with IL-1β, and this treatment produced a similar sparse distribution of clots within collectors. Activation of G-protein coupled receptor PAR-1 with thrombin acts synergistically with TNFα on TF expression on blood endothelium(Liu,& al,2004) and that treatment has the potential to augment the effect of cytokines. Surprisingly, injection of 2.5 IU thrombin alone had no effect on intralymphatic coagulation away from the site of injection but as expected led to massive coagulation within blood vessels at the injection site (**Supplementary Fig. 2**). Next, we wanted to block lymphatic thrombomodulin by injection of chemically inactivated thrombin. Using a covalent inhibitor phenylmethanesulfonyl fluoride (PMSF), we found that only 80% of the enzyme (six-times times longer clotting time as compared to thrombin^ac^) was inactivated. Recurrent intradermal injection of partially inactivated thrombin (thromb^in/ac^) had an effect similar to cytokines treatment and induced the formation of intralymphatic clots within specific locations of lymphatic collectors but without the bystander coagulation within blood vessels **(Fig 1C and Supplementary Fig. 3)**. Even though fibrin was deposited only in a few collector branches (but always in at least one collector segment in all 9 tested ears), few lymph clots were sufficient to occlude lymphatic drainage from all lymphatics efferent to that collector **(Fig. 1D)**. There was no cumulative effect of combined treatment with cytokines and thrombin on lymph clotting and like with the injection of individual factors only discrete locations in collectors were functionally occluded with clots. Lymphatic occlusion induced by the partially inactivated thrombin (thrombin^in/ac^, **Fig. 2A**) or a mix of TNFα and Il-1β cytokines and thrombin^in/ac^ (**Fig. 2B**) could be prevented with TF blocking antibodies. Confocal images of fibrin-occluded lymphatics stained for TF revealed that lymph clotting co-localized with high expression of TF on lymphatic endothelium. Heterogeneity of TF expression after stimulation with TNF or partially inactivated thrombin resembles the similar discontinuous pattern of CCL21 expression previously reported by our group (Kilarski,& al,2013). It is also characteristic for blood vessels where TF expression on the endothelial cell varied strongly among TNF-alpha-stimulated endothelial cells (Kirchhofer,& al,1994) and it is localized mostly at the vascular bifurcations (Lupu,& al,2005, van Hinsbergh,2012). Dendritic cells (DC) were absent from intralymphatic cloths in triple-factor stimulated lymphatics but filled the lymphatic clots along the zone of high TF expression in thrombin^in/ac^ treated skin (**Fig. 2B**).

**Figure 2.**
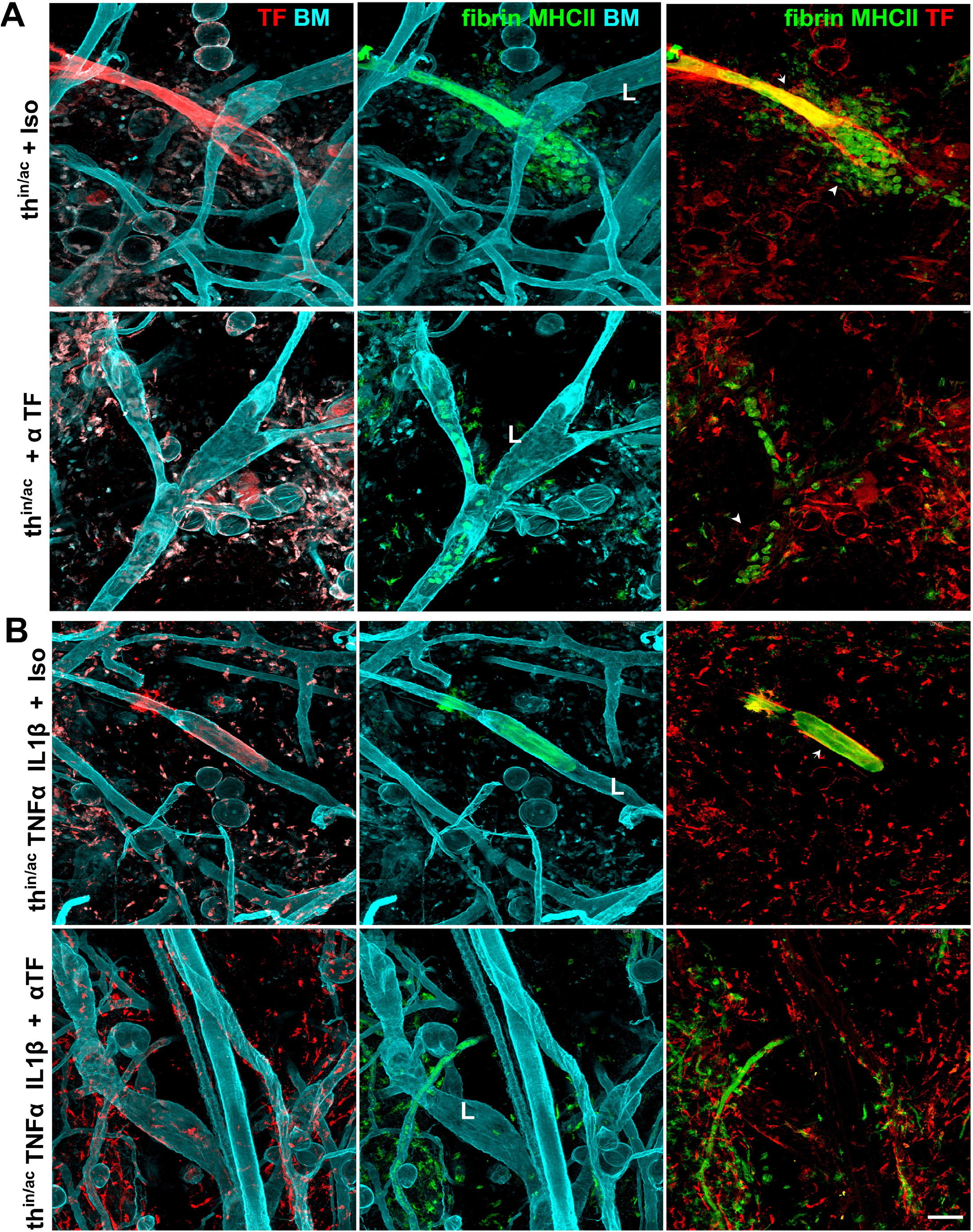
Intralymphatic coagulation induced by cytokine or partially-inactivated thrombin is dependent on tissue factor expression on lymphatic endothelium. Confocal whole-mount images of lymphatic collectors occluded by intralymphatic clots after two consecutive intradermal injections with pro-clotting factors with isotype control or tissue factor blocking antibody (21E10) Basement membrane (BM) was stained with collagen IV, antigen presenting cells are stained with MHCII. **A**. Injection of partially inactivated (thrombin^in/ac^) together with isotype control antibody induced high expression of TF on discrete location of collectors. Locations of lymph clotting coincided with high expression of tissue factor (TF) on collector endothelium (red within collector’s vessels). Collectors that express a high level of TF were densely infiltrated with MHCII-positive antigen presenting cells. TF-blocking antibody (αTF) completely abolished clot formation but not clustering of antigen presenting cells within lymphatic collectors. **B**. Similar, in the presence of the isotype control antibody, mix of partially inactivated thrombin (thrombin^in/ac^) and cytokines (TNFα and IL1β) induced fibrin clot formation in discrete locations within downstream collecting vasculature. In contrast to thrombin^in/ ac^ alone, these clots were not infiltrated with MHCII-positive leukocytes. When αTF antibody was injected with the pro-clotting mix, lymph clots did not form within lymphatic collectors, and TF expression was reduced or absent on lymphatic endothelial cells. Arrows point to fibrin clots, arrowheads, MHCII-positive antigen presenting cells. L marks lymphatic collectors. TF intensity was normalized to the TF expression level of interstitial cells. Note that lymph clots are autofluorescent in the red channel due to cross-reactivity of anti-rat antibody with clot-entrapped mouse IgGs, hence the TF signal can be appreciated only at the edges or outside the clot. Scale bar: 50μm.

### Lymphatics neutralize blood-born thrombin and preserve lymph drainage

These experiments suggested that the lymphatic anti-clotting mechanism is primarily focused on the inhibition of the idling generation of thrombin or self-activated thrombin formed during the amplification phase of the clotting. Even in hemostasis, there is a continuous TF-dependent activation or ‘idling’ of the clotting system that results in a constant, low-level formation of active thrombin. The minimal level of the active factor VIIa that co-factor TF is necessary for the activation of blood clotting after tissue injury (Jesty,& al,2005, Mackman,2009). The idling mechanism of activation of blood-borne thrombin, in addition to the dual-mode of activation of thrombin receptors, could explain intricate responses we observed after injection of active (thrombin^ac^), partially active (thrombin^ac/in^), or completely inactivated thrombin (thrombin^in^). Thrombin has two types of receptors, thrombomodulin and a protease-activated G-coupled receptors (PARs)(van Hinsbergh,2012), with thrombomodulin and PAR-1 existing mainly on endothelial cells. PAR-1 must be activated by thrombin proteolytic cleavage, hence inactivated thrombin cannot initiate PAR-1 signaling and thrombin responses in cells. On the contrary, thrombin binding by thrombomodulin, the cofactor that alters thrombin specificity from fibrinogen to protein C, is independent of the thrombin enzymatic state (Esmon,1995). Therefore thrombomodulin can bind thrombin when its enzymatic activity is inhibited by small molecular inhibitors (Fuentes-Prior,& al,2000). Thrombomodulin-bound active thrombin shifts its enzymatic specificity from fibrinogen to protein C, which proteolytic activation, in turn, leads to the destruction of co-factors V and VIII and inhibition of clotting process (Conway,2012). Hence, blocking thrombin binding sites on thrombomodulin with inactive thrombin should specifically inhibit the activation of PARs receptors on variety of stromal cell types that normally lead to activation of inflammatory response and compensatory anti-clotting pathways. Treatment of thrombin with the potent, small-molecule covalent inhibitor (p-amidonophenyl)methane sulfonyl fluoride (APMSF) inactivates 99.75% of the enzyme (Laura,& al,1980) and prolongs the clotting time by 400-times as compared to thrombin^ac^. Consequently, injection of PMSF-treated and APMSF-treated thrombin initiated vastly different responses in the skin (**Fig. 3A**). As expected, intradermal injections of saline had no effect on intralymphatic clotting, dextran drainage **(Fig. 3B and C, left)**, or tissue plasminogen activator (tPA) expression in lymphatics **(Fig 3D, left)**. Injection of fully active thrombin (thrombin^ac^), induced local coagulation within blood vessels but had no effect on lymph clotting in remote lymphatic collectors **(Fig. 3B and 3C, middle)**. Additionally, at the site of injection thrombin^ac^ induced robust expression of tPA in various tissue cells, i.e., adipocytes, macrophages, and blood vessels but only at the site of injection **(Fig 3D, middle)**. On the other hand, APMSF-inactivated thrombin (thrombin^in^) led to rapid and massive lymph clotting in most segments of draining lymphatic collectors **(Fig. 3B and C, right)**. In thrombin^in^ injected skin, intra-lymphatic clots were surrounded by increased levels of tPA within collectors standing out in the background of the other tissue structures **(Fig. 3D, right)**.

**Figure 3.**
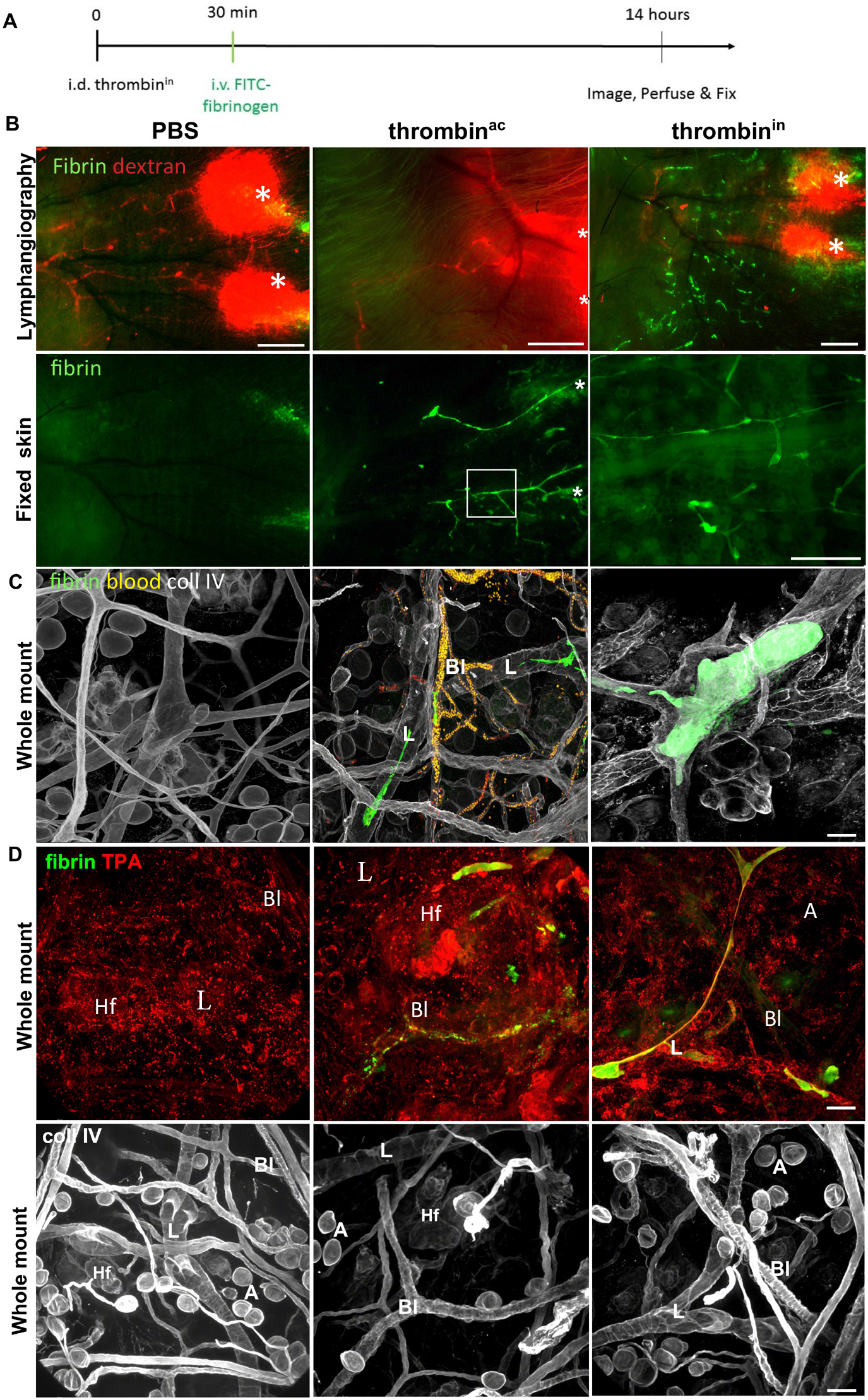
Completely inactivated thrombin blocks lymphatic drainage. **A**. Schematics of experimental design. Intradermal injection of saline, active thrombin (thrombin^ac^) or completely inactivated thrombin (thrombin^in^) was followed by intravenous injection of fluorescently labeled fibrinogen. The mouse is fixed by perfusion 13 hours later. **B. Top**: Lymphangiography and intra-lymphatic fibrin in collectors of live tissue 5 hours after intradermal injection. **Bottom**. Imaging of fluorescent insoluble fibrin clot in fixed-perfused dorsal skin flap after cartilage and ventral skin were removed. Two adjacent injections of 2.5 μl PBS had no effect on the collector-drainage later (**PBS, top**). Fibrin formed only in the interstitium at the injection sites (**PBS, bottom**). Active thrombin (2.5 IU; thrombin^ac^) increased hyperpermeability of collectors but did not block their drainage (**Top, thrombin**^**ac**^). Fibrin formed within blood vessels around the site of injection (**Bottom, thrombin**^**ac**^). In contrast to thrombin^ac^, intradermal injection of inactive thrombin (thrombin^in^) occluded remote lymphatic collector vessels (**Top, thrombin**^**in**^). Drainage discontinuities visible on lymphatic lymphangiography are bridged by green fibrin deposits, indicating an incomplete occlusion. In contrast to continuous fibrin clots induced in blood vessels by thrombin^ac^, thrombin^in^ induced the formation of discontinuous clots in a large number of remote lymphatic collectors but had no effect on plasma coagulation in blood vessels (**Bottom thrombin**^**in**^). **C**. Higher magnification confocal images 5 hours after the second injection of compounds indicated in B. Collagen IV stained lymphatic collectors (L) and blood vessels (Bl) identified by their characteristic morphology (valves, uneven diameter along the vessel length). In PBS injected mice, fibrin clots were not formed away from the injection site. CD31 showed normal cell-cell junctions between lymphatics (**PBS**). Active thrombin blocked blood circulation, trapping autofluorescent erythrocytes (yellow) within blood vessels (**Bl**) around the injection site. Rare intralymphatic clots (**L**) could be found only around the thrombin^ac^ injection site (**Thrombin**^**ac**^). In the **Thrombin**^**in**^ injected skin, discontinued fibrin clots in remote lymphatic collectors were observed. **D. Top**. 16 hours after injection of **PBS**, tissue plasminogen activator (TPA) is randomly distributed within the skin but is also stain morphologically distinct macrophages (M)., erythrocytes trapped within blood vessels, TPA-positive macrophages but also large vessels were seen around the site of **Thrombin**^**ac**^ injection. In the skin injected with **Thrombin**^**in**^, TPA strongly labeled collecting lymphatic vessels filled with residual, centrally-located fibrin clots. **Bottom**. Imaging of the same field stained with collagen IV depicted tissue stationary structures, i.e., blood vessels, lymphatics, and adipocytes. A-adipocytes, L-lymphatic collectors, Bl-blood vessels, M-macrophages, Hf-hair follicle. The intensity of TPA was normalized to its signal level in interstitial cells. The intensity of lymphatic fibrin deposits was normalized to signal level of clots at the sites of injection (*). Scale bar: B–500 μm, C-50μm.

The patterns of thrombosis in the dermis injected with different formulations of thrombin correlated with different kinetics of clot fibrinolysis. Saline injections led to the formation of dermal clots at the site of the needle insertion but had no effect on blood or lymphatic vessels thrombosis **(Fig. 4A, top)**. Thrombin^ac^ repeatedly occluded local blood circulation but not lymphatics around the sites of injections and these blood thrombi were almost entirely cleared from the blood vessels within hours **(Fig. 4A, center)**. Partially inactive thrombin (thrombin^in/ac^) induced clots formed within the scattered location of lymphatics only after the second injection suggesting that the initial injection of thrombin^in/ac^ stimulated scattered acquisition of a pro-thrombotic phenotype within collectors (**Fig. 4A, bottom**). Contrary, the completely inactive thrombin (thrombin^in^) induced massive fibrin clotting in most draining collectors already 2 hours after the first injection. Like blood thrombi induced by thrombin^ac^, lymphatic clots induced by thrombin^in^ were mostly cleared from collectors within a day after the injection **(Fig. 4B, top)**.

**Figure 4.**
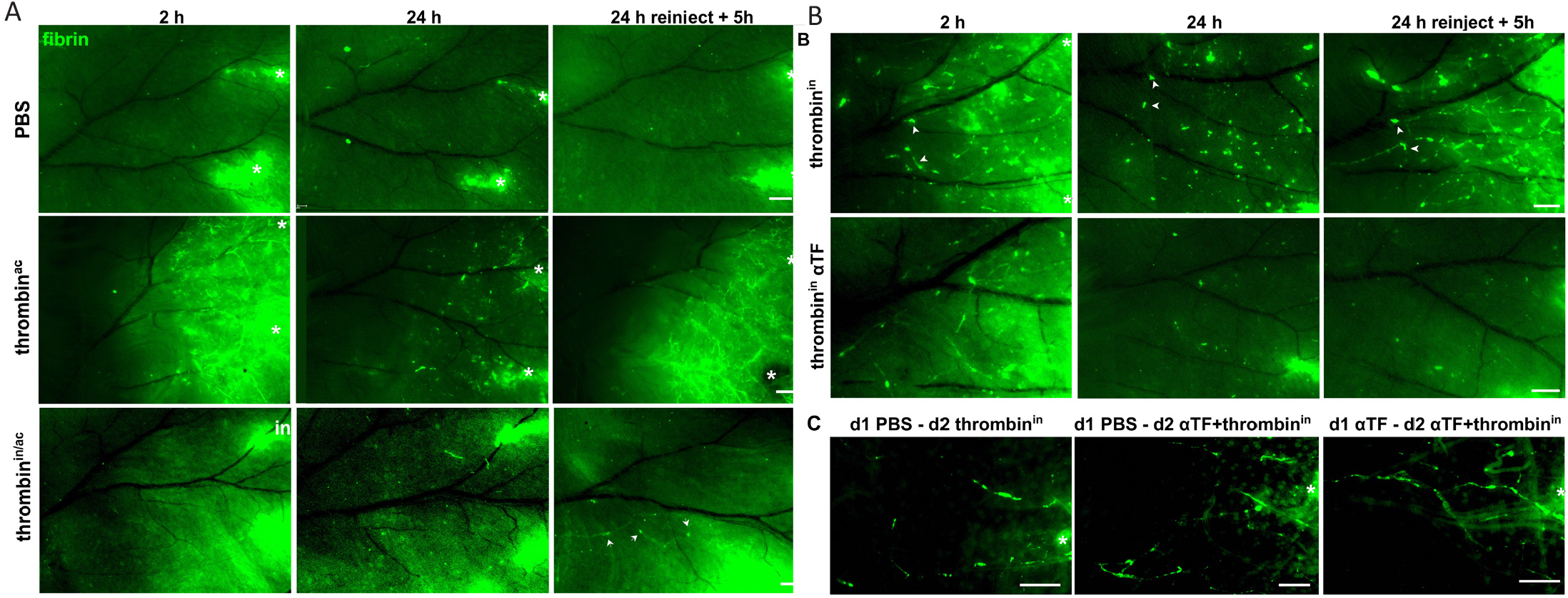
Completely inactivated thrombin instantly but transiently occlude lymphatic collectors by a tissue factor-independent mechanism. Intravital imaging of fluorescent fibrin clot formation within the ear skin vasculature. Fluorescent fibrinogen was administered i.v. immediately after 2.5 μl intradermal injection of indicated compounds in two adjacent injection sites on a single ear dorsal dermis. In all images, lymphatics drain from the right (distal) to the left (proximal). The functional blood vessels appear dark in all images as tissue was fixed by perfusion to clear the vessels from non-coagulated fibrinogen. **A**. Top: Injections of PBS result in injury of blood vessels, plasma leakage, and clot formation within the dermis, which was restricted to immediate surroundings of injection spots. **Center**: Injections of active thrombin (thrombin^ac^, 2.5 IU) resulted in coagulation in blood vessels, blood vessel leakiness, and fibrin clotting within the dermis in the area of the ear where injection edema formed. Intravascular and interstitial clots were largely cleared within 24 hours after the injection, however, they could be re-created with another intradermal injection of active thrombin^ac^. **Bottom**: Similar to PBS injections, a single dose of partially inactivated thrombin (thrombin^in/ac^) caused leakage of plasma and intradermal fibrin clotting around the site of injection but had no effect on blood or lymphatic vasculature. However, reinjection of thrombin^in/ac^ 24 hours later induced the formation of intralymphatic clots at discrete lymphatic locations, downstream to the sites of injection. Arrows point to single intralymphatic clots defined by their sharp edges, in contrast to the whole lymphatic that is visible due to non-coagulated fibrinogen labeled lymph. **B**. Ears after injection with thrombin^in^ inactivated with APMSF (400-fold decrease in activity, thrombin^in^). **Top**: Single dose of thrombin^in^ (co-injected with irrelevant rat antibodies) resulted in rapid formation of fibrin clots within most draining lymphatic collectors, which were mostly cleared 22 hours later. Reinjection of thrombin^in^ re-created clotting pattern within lymphatics. Arrowheads point the same intralymphatic clots visible at 2 and 24 hours after first injection and 5 hours after the second injection. **Bottom:** Co-injection of tissue factor-blocking (αTF) antibodies and thrombin^in^ did not inhibit instant intralymphatic clot formation, however reinjection of αTF and thrombin^in^ 24 hours later did not induce the formation of intralymphatic clots. **C**. Neither pre-injections with PBS (injection injury) nor αTF antibody (day 1) inhibited intralymphatic clot formation induced with thrombin^in^ and αTF co-injection on day 2 and at 24 hours after the first injection. Images were taken from ears fixed 5 hours after the second injection. The intensity of lymphatic fibrin deposits was normalized to the signal level of clots at the sites of injection (*). Scale bar: 250μm.

### Initial intralymphatic clotting is TF-independent

Fibrinolysis dependents also on the feedback from the thrombin activity. However, the dynamic of intralymphatic clotting/fibrinolysis differed significantly from thrombotic/ thrombolysis processes characteristic for blood vessels (2019). The first co-injection of TF-blocking antibodies with thrombin^in^ had no effect on intralymphatic clotting **(Fig. 4B, bottom)**. However, re-injection later of TF-blocking antibodies with thrombin^in^ 22 hours completely blocked thrombin^in^-dependent lymph clot formation within collectors **(Fig. 4B, right)**.

This is confirmed by the fact that neither the injection itself with saline nor anti-TF antibodies prevented the formation of intralymphatic clots blocked the formation of intralymphatic clots **(Fig. 4C**), confirming that blocking the anti-thrombotic activity of lymphatics led to stimulation of lymphatic pro-thrombotic activity after recurrent and indirect stimulation of lymphatic endothelial cells with inactive thrombin. it seems likely that initial activation of the clotting pathway leads to sensitization to the extravascular TF by shifting the intralymphatic hemostatic balance to fibrinolytic, where activated endothelial cells upregulate the production of tPA **(Fig. 3D)**. Injection with TF-blocking antibodies and subsequent injection of skin with TF-blocking antibodies and thrombin^in^ did not inhibit lymph clotting **(Fig. 4C)**. Instead, idle extravascular TF activity (the idling extrinsic pathway) or contact activation (intrinsic) pathway initially sufficed for the accumulative lymph clotting and become insufficient in the presence of high levels of plasmin.

### Sterile inflammation in collectors results in lymph clotts that blocks lymphatic drainage

Necrosis of endothelial cells within lymphatic collectors leads to localized sterile inflammation that should have multifactorial pro-hemostatic effects on the transported lymph(Chen,& al,2010). First, the elimination of thrombomodulin anti-clotting mechanism inherently linked to lymphatic endothelial cells promotes amplification of contact-dependent pathway on exposed nucleic acids and polyphosphates from dying cells(Esmon,2010). Also, decellularization of the lymphatic surface should prolong the occlusion of lymphatics as it eliminates the source of endothelium-derived tissue plasminogen activator(Wu,& al,1996) that is a cause of the fast clearance of intralymphatic clots from TF-expressing or thrombomodulin-blocked lymphatics **(Fig. 4B and 4C)**. Finally, sterile inflammation leads to infiltration of TF-expressing activated monocytes that could directly induce lymph clotting by extrinsic pathway(Egorina,& al,2008).

To induce sterile inflammation of collecting vessel endothelium we choose agents that have a direct toxic effect on living cells, a cell lysing detergent Triton-X-100 (Triton) and FeCl_3_, a catalyst generating hydroxyl and hydroperoxyl radicals from mitochondria-derived hydrogen peroxide and it is used in TF-independent blood vessel thrombus formation models(Wang,& al,2005). Intradermally injected dextran mixed with 1% detergent Triton-X-100 was drained initially by collecting lymphatics, to be leaked out of the lymphatic collector into the remote tissue only minutes later **(Fig. 5A and 5B)**. Lymphangiography repeated on the next day showed that draining of cell lysis agent blocked the tissue drainage completely and instead lymphatic collectors were filled with continuous lymph clotting **(Fig. 5C)**. As expected, angiography performed day after intradermal injection of Triton showed that blood circulation was lost around the injection spots, confirming Triton in vivo universal toxicity and capability of blocking circulation. Optical sectioning of whole-mount preparation of the injected skin revealed that fibrin clot filled the completely lumen of the draining lymphatic, which lost the fine organization of cell-cell junctions **(Fig. 5D)**. However, the Triton toxicity was limited to the collectors specifically draining detergent as afferent lymphatic draining control region but joined to the fibrin occluded collector vessels, but also blood capillaries surrounding occluded lymphatics, preserved their cellular organization. Already one day after the treatment surrounding tissue and fibrin-occluded lymphatics were populated by rounded inflammatory cells. Different mechanism of FeCl_3_ toxicity that is dependent on free radical generation from H_2_O_2_ produced in mitochondria surprisingly lead to the same final effect, with blocked collector drainage and deposition of fibrin clots within lymphatic **(Fig. 5E**). In addition, similar to Triton X-100, intradermal injection of ferric chloride resulted in complete cessation of blood circulation around dermis injection sites. Fibrin clots filled the complete lumen of lymphatics draining ferric chloride, which also lost its fine CD31-delinighthed junctional organization **(Fig. 5F)**. Likewise, Triton x-100, FeCl_3_ toxicity was limited to draining lymphatic as lymphatics directly connected to a fibrin-occluded collector but draining lateral skin (internal control) and surrounding blood capillaries preserved their cellular arrangement. Also, rounded inflammatory (CD45+) cells infiltrated the draining region and crowded within the fibrin of occluded lymphatics.

**Figure 5.**
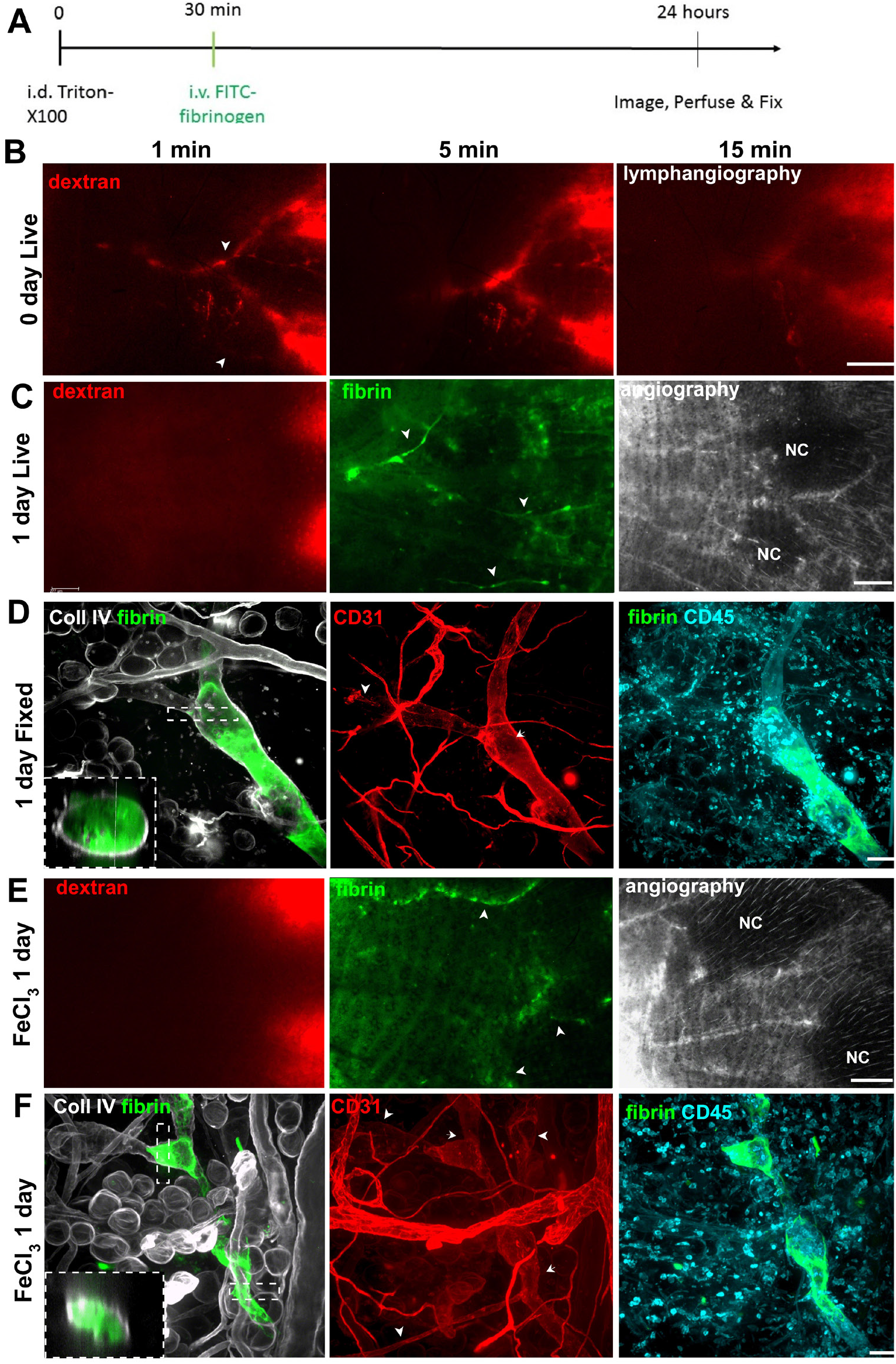
Sterile inflammation induced by Triton or FeCl_3_ injure draining lymphatics and cause the formation of lymph clots that completely occlude collector drainage. **A**. Schematics of experimental design. Intradermal injection of 10% Triton-X-100 or FeCl_3_ was followed by intravenous injection of fluorescently labeled fibrinogen, and mouse was fixed/perfused at 24 hours. **B-D**. Fluorescent dextran mixed with Triton-X-100 (TX100) was injected in the distal ear skin in two radially distinct locations that are drained by separate branches lymphatic collectors. **B**. Intravital microscopy of early time points after injection of dextran-TX100. From **Left** to **Right:** marking the collectors at 1 minute; 5 minutes after injection, leakage from the collecting vessels is visible; 10 minutes later, the extent of dextran diffusion from the leaky collectors mask their original drainage path. **C**. 24 hours after a mix of dextran-TX100 was injected, fluorescent dextran tracer re-injected i.d. is not drained by lymphatics. The chase with FITC-labeled fibrinogen leaked into the ear dermis and polymerized in the tissue, indicating disperse tissue injury but also within lymphatic collectors that were initially been draining the dextran-TX100 mix. Arrowheads in B and C mark the same collector locations. Fluorescent angiography performed 24 hours after i.d. injection revealed blood circulation deprived zones (**NC**-no circulation zones) in the skin where dextran-TX100 mix was injected. **D**. The whole-mount imagining of the skin collecting lymphatics fixed-perfused one day after the TX100 treatment. **Left:** Fluorescent fibrinogen that leaked from blood vessels polymerized within draining lymphatics, completely occluding their lumen. Insert represents a 5μm transverse optical section through an occluded lymphatic collector. **Middle**. The intracellular localization of CD31 is present in lymphatic draining control lateral ear side stains cell-cell junctions before the valve emptying into the clot-occluded lymphatic collector (arrowhead). In contrast, junctional staining is lost in the TX100-damaged collector, and instead, CD31 signal is evenly distributed across the portion of the collector where fibrin clot formed (arrow). **Right:** CD45 staining demonstrates a swarm of neutrophil-like immune infiltrates in the interstitial tissue. **E, F**. The study at one day after intradermal injection of FeCl_3_, demonstrate similar phenotypes as with dextran-TX100. **E**. Intravital microscopy. **Left:** the perfusion of collectors draining the peripheral region of dorsal ear skin was blocked as demonstrated with lymphangiography performed by i.d. dextran injection as in B. **Middle**: Fibrin formed in the dermis and marked lymphatic collecting vessels (arrowhead). **Right:** Similar to TX100 injections, i.d. administration of ferrous ions was toxic to the skin, leading to blockage of blood circulation around the sites of intradermal injection (no circulation zones, **NC**) as revealed by fluorescence angiography. **F**. The whole-mount imagining of the skin. **Left:** Fluorescent fibrinogen that leaked from blood vessels polymerized within draining lymphatics and at certain locations completely occluding their lumen. Insert represents a 5 μm transverse optical section through an occluded lymphatic collector. **Middle:** CD31 staining in healthy lymphatic collector draining lateral control region mark cell-cell junctions and a valve (arrowheads). In contrast, in medial collectors that drained ferrous ions, junctional staining is lost and CD31 signal dispersed within the fibrin clots (arrows). **Right:** Fibrin-occluded lymphatics and surrounding dermis were infiltrated with inflammatory cells 24 hours after the treatment. The intensity of CD31 was normalized to its signal level in the lymphatics draining control region. Scale bar: B, C, E-500μm; D, F-50μm.

### Decellularization of collectors leads to complete and long-lasting lymphatic occlusion

Injection of general toxins envisaged the importance of anti-clotting properties of the endothelium from lymphatic collectors. It also pictures the ease and specificity with which they can be targeted, additionally proving that blocking of only a few lymphatic collectors ceases the drainage from the whole lymphatic tree. Including intact vessels draining the lateral side of the ear skin which are connected to the occluded lymphatic (**Fig. 5D and 5F**). This approach unspecifically exerts bystander toxicity to any cell type present at the site of injection. Also, toxins cannot be inactivated, instead, they can freely diffuse outside the collectors potentially injuring cells in remote locations. Instead, photodynamic therapy (PDT) offers a solution to all these limitations. PDT is a combined physicochemical treatment were intradermally injected photosensitizer acts as proto-toxin, which is activated and destroyed with laser light at a defined location. This treatment assures that the toxicity of free radicals diffusing over short distances (up to 7 μm) will be strictly limited to collectors’ associated cells(Kilarski,& al,2014). Large, 100nm in diameter Visudyne® liposome enclose an active compound, a small molecular verteporfin, therefore their diffusion potential in the tissue was minimal. Practically, Visudyne® liposomes were either directly injected into the lymphatic collectors or retain in the skin interstitium. To avoid direct toxicity to the dermis that could result in enhanced trafficking of immune cells from the afferent injury site, the injection spots were painted black and covered with foil immediately after the injection only leaving the exposed collectors. We reached 98% of the reproducibility of the anti-lymphatic treatment by confirming the phototoxic self-annihilation of Visudyne® within lymphatic collectors. This was done by imaging Visudyne® fluorescence before and after irradiation, which allowed us to eliminate mice with incompletely or poorly injected collectors. The routine anti-lymphatic treatment occluded all collectors draining peripheral dorsal skin within 24 hours after PDT with continuous lymph clots **(Fig. 6B**). Likewise, injection of directly-acting toxins, Triton or ferric chloride (**Fig. 5**), and in contrast to TF-stimulation **(Fig. 1-2)** or thrombomodulin blocking **(Fig. 3-4**), PDT always resulted in the formation of continuous intralymphatic fibrin clots in all vessels that drained photosensitizer **(Fig. 6B)**. In contrast to directly-acting toxins, PDT resulted in much lower collateral damage to the tissue at the injection site that was manifested in minimal leakage of fluorescent plasma into a tissue (compare **Fig 5C and D with Fig. 6B**). Endothelial cell death deduced from degeneration of cellular junctions was specific to endothelial lining coming in direct contact with a photosensitizer, leaving afferent vessels draining lateral side of ear dermis intact **(Fig. 6D)**. Basement membrane that supports endothelium and muscle cells of lymphatic collectors **(Fig. 6D)**, remained as a scaffold where lymphatic clots formed occluding the lumen of “ghost vessels” **(Fig. 6D)**, named after similar blood vessel-derived structures that were described earlier by Inai(2004). Even though the basement membrane of collecting lymphatics remained intact, some degeneration occurred as BM particles could be found trapped within the fibrin clots **(Fig. 6D, insert)**. Fibrin-occluded lymphatics were densely infiltrated by rounded neutrophil-like immune cells from day two after PDT. Leukocyte clusters persisted at the cone of advancing endothelial cells that regenerated ghost vessels **(Supplementary Fig. 4)**, possibly providing directional and stimulating cues by the release of, e.g. TGFβ **(Fig. 6E)** to re-growing endothelium (van Meeteren,& al,2012). In contrast to fibrin deposits within ghost vessels, which began to clear from day 4 onward, massive leukocyte infiltration persisted throughout the regeneration period of lymphatics even though **(Fig. 6E)**. Fibrin deposits were eventually either cleared or replaced with abnormal non-sheet-like masses of basement membrane matrix, containing laminin **(Fig. 6F)**, collagen IV, collagen III and tenascin C. Basement membrane deposits eventually evanesced from vessels approximately by day 9, which coincides with lymphatics re-gaining their drainage functionality ((Kilarski,& al,2014) and **Fig. 10b**). Interestingly, similar intralymphatic matrix deposits, composed of collagen type I and IV we found within peritumoral collecting lymphatics in implanted dermal B16-F10 melanoma, which was paralleled by the abnormal, discontinued organization of lymphatic endothelial cells **(Supplementary Fig. 1)**.

**Figure 6.**
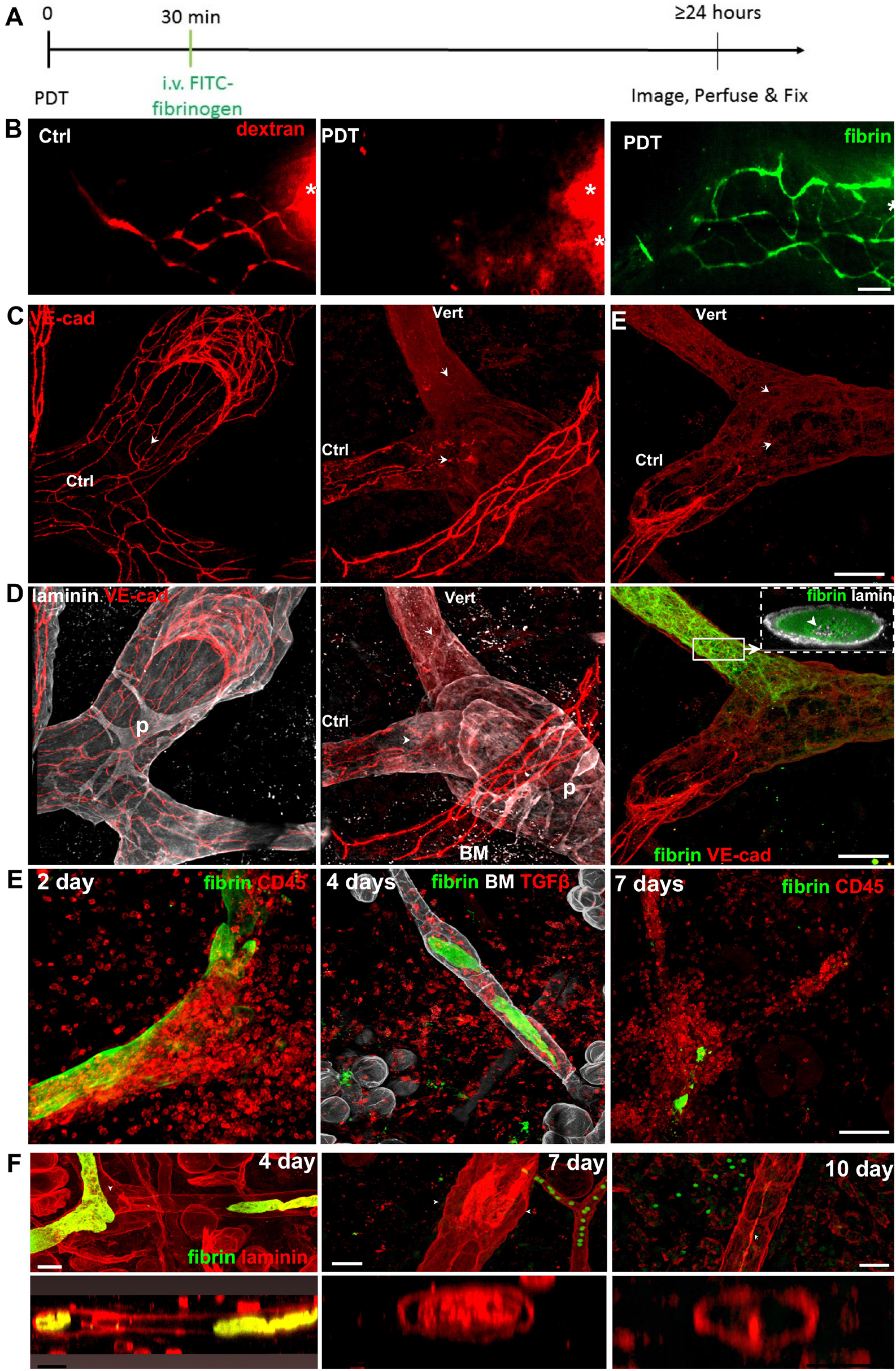
Specific de-cellularization of lymphatic collectors leads to complete and long-lasting lumen occlusion. **A**. Schematics of experimental design. Endothgelium was stripped from lymphatic collectors with a photodynamic therapy (PDT). Specifically, intradermal (i.d.) injection of photosensitizer was followed by collector imaging (verteporfin lymphangiography) and subsequent irradiation of dorsal ear skin (PDT). 20-30 min after PDT, FITC-fibrinogen was injected intravenously, and ear lymphatics were imaged the following day. The mouse was fixed/ perfused on the indicated day for whole mount imaging. **B**. Intravital microscopy, C-F-confocal imaging of whole-mount skin preparations. **B. Left:** Saline injection and repeated lymphangiography had no effect on collector perfusion in the control ear. **Middle:** PDT leads to occlusion of lymphatic collectors, which stop draining i.d. injected fluorescent dextran. **Right:** The same collector networks become occluded with insoluble fibrin that leaked from blood vessels. **C-E**. Whole-mount imaging. **Left:** VE-cadherin is localized at cell-cell-junctions in regular lymphatic collectors of the ear skin. **Middle, Right:** VE-cadherin organization is lost in vessels that drained photosensitizer, while in the same ear, control lymphatic (**Ctrl**) draining lateral to the Visudyne® draining collectors or neighboring blood vessel (5 μm away) were not affected by irradiation. **D. Left:** Lymphatic endothelial cells (**VE-Cad**-VE-cadherin) were enclosed within the basement membrane (stained for laminin). P-basement membrane imprints of collecting vessel pericytes. **Middle:** Morphologically unaffected, cell-deprived basement membrane tubes- ‘ghost vessels’- one day after PDT. BM scaffold was preserved in PDT-decellularized collectors. **Right:** Ghost vessels were filled with fibrin clots that blocked the whole lymphatic lumen. Insert shows 5μm cross-section of the fibrin-occluded collector. Frequently, basement membrane particles could be found trapped within a clot (arrowhead). **E**. In addition to intralymphatic lymph clotting, ghost vessels become infiltrated with TGFβ^+^ leukocytes that densely pack within the collector. **F**. From day 4 to 7 fibrin clots are cleared and partially replaced with tissue deposits containing basement membrane elements of laminin that are subsequently removed from vessel lumen(day 7 to day 10). The intensity of CD31 was normalized to its signal level in the lymphatics draining control region. The intensity of lymphatic fibrin deposits was normalized to the signal level of clots at the sites of injection (*). Scale bar: B, 250μm, C, D, E-100μm, F-25μm.

### Endothelial cell death precedes tissue factor-independent lymph clotting

We could not find even a theoretical mechanism explaining why the killing of lymphatic endothelial cells but not its basement membrane scaffold should lead to occlusion of collectors without the implication of a secondary mechanism, namely lymph clotting. This also suggests that the primary function of lymphatic collectors could be the control of the lymph fluid state.

Live imaging of skin collector vessel endothelium expressing Tomato-fluorescent protein under Prox1 promotor, with basement membrane stained for collagen IV showed that irradiated cells draining proto-toxin Visudyne® died between 1 and 3 hours after PDT (**Video 1, Fig. 7A**). Vertical afferent lymphatic collector served as the internal control for imaging-dependent bleaching and phototoxicity and showed that only prior laser irradiation of drained photosensitizer but not subsequent fluorescence imaging evokes the death of lymphatic endothelium. As expected, there was no quenching of intracellular Tomato proteins during imaging with a low-power mercury lamp used in our fluorescence imaging setup(Guc,& al,2014).

**Figure 7.**
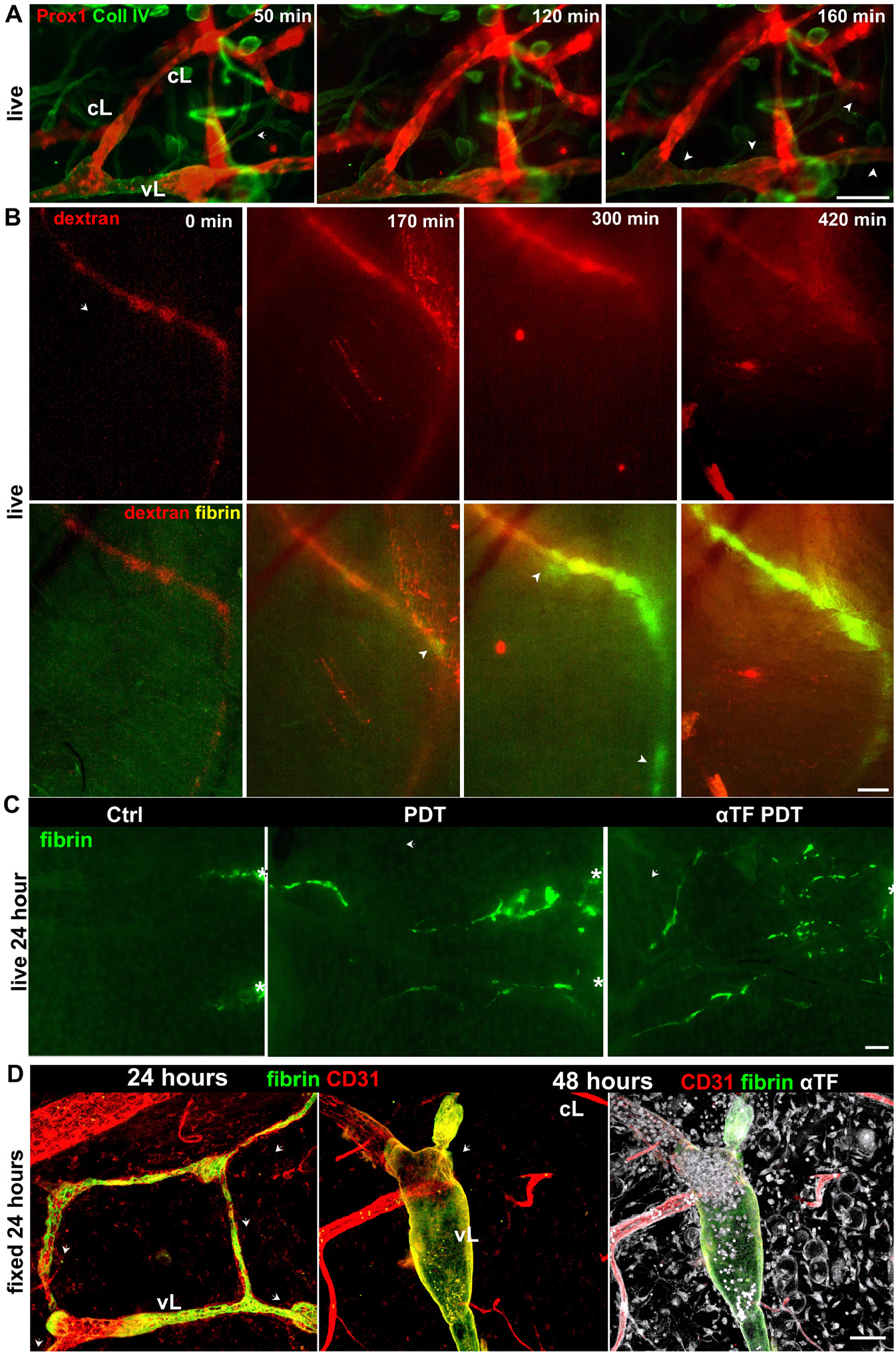
Lymph clotting and drainage blockage is preceded by lymphatic cell death. **A**. Time course of lymphatic collector cell death after PDT. Intravital imaging of the exposed dorsal ear skin of Prox1-Tomato mouse that was live-stained for basement membrane with collagen IV immediately after photodynamic therapy (PDT). Single lymphatic collector drained Visudine® and was irradiated (**vL**), while two lateral lymphatics were irradiated but did not drained photosensitizer and served as an internal control for imaging-related bleaching and phototoxicity (**cL**). Arrowheads indicate locations of collecting lymphatic where endothelial cells were undergoing necrosis. **B**. Time course of intravascular lymph clotting and drainage block in lymphatics after PDT. Lymph clots start forming 170 minutes after decellularization of collectors and is completed 300-420 minutes (5-7 hours) later. Lymphatic drainage ceases after 5-7 hours after lymphatic PDT. Arrowheads point to fibrin deposits (green) within collector’s vessel initially draining fluorescent dextran (red). Fibrin forms during the first day (day 0) after lymphatic decellularization. **C**. Fluorescent fibrin clot imaging, intravital microscopy. Mechanism of post-PDT intralymphatic clotting was analyzed by blocking the extrinsic pathway with tissue factor (TF) blocking antibody (αTF) injected together with the photosensitizer and 1 hour after PDT. TF blocking had no effect on intravascular lymph clotting. **D**. Confocal microscopy. **Left**. Despite the presence of αTF, fibrin formed within lymphatics 24 hours after PDT. **Middle**. Presence of TF-blocking antibodies had no effect on intralymphatic clot stability 48 hours after PDT. **Right**. Blocking the TF pathway had no effect on TF-positive leukocyte infiltration into decellularized lymphatics after PDT. A-D. The intensity of lymphatic fibrin deposits was normalized to the signal level of clots at the sites of injection (*). Arrows indicate the direction of Visudyne® flow. Scale bar: A-C-100 μm, D-50 μm.

Lymphatics expressed Tomato fluorescent protein in their cytoplasm that either should diffuse out of the necrotic cell or be compartmentalized into fluorescent apoptotic or autophagosomes bodies during programmed cell death as it is shown in **Video 2**. We showed before that cell killing with PDT occurs by necrosis as cells become permeable to propidium iodide within 1.5 hours after the treatment ((Kilarski,& al,2014) also **Video 3**). Here we show that most endothelial cells treated with PDT diffuse their cytoplasmic proteins, the more informative indicative of the final stage of necrotic cell death. Despite that collector’s endothelium dies off within 2.5 hours after PDT vessel occlusion started no earlier than 3 hours after PDT **(Fig. 7B)**, which is in agreement with our previous results(Kilarski,& al,2014). The cessation of dextran drainage between 3 and 5 hours after PDT correlated with increased deposition of fibrin within collectors that reached its maximum 7 hours after PDT. As compared to blood vessel occlusion, the process that is completed within minutes after the injury, occlusion of lymphatic collector after PDT-induced decellularization was a slow process that took at least an hour and began with fibrin deposition on the wall of decellularized lymphatic and continued toward the center of the vessels **(Video 4)**. In contrast to lymph clotting provoked by cytokines-induced TF-expression, intralymphatic clotting after PDT-decellularization could not be prevented with TF blocking antibodies **(Fig. 7C-D)**. Also, TF-blocking did not inhibit the infiltration of leukocytes into decellularized lymphatics, despite the fact that antibody was present in the tissue two days after the treatment. Most importantly, we confirmed our previous observation that there is a temporal gap between endothelial cell death and lymphatic occlusion, which indicated that at least during this lag period decellularized lymphatics were able to drain fluid without the presence of endothelial cells.

**Video 1.**
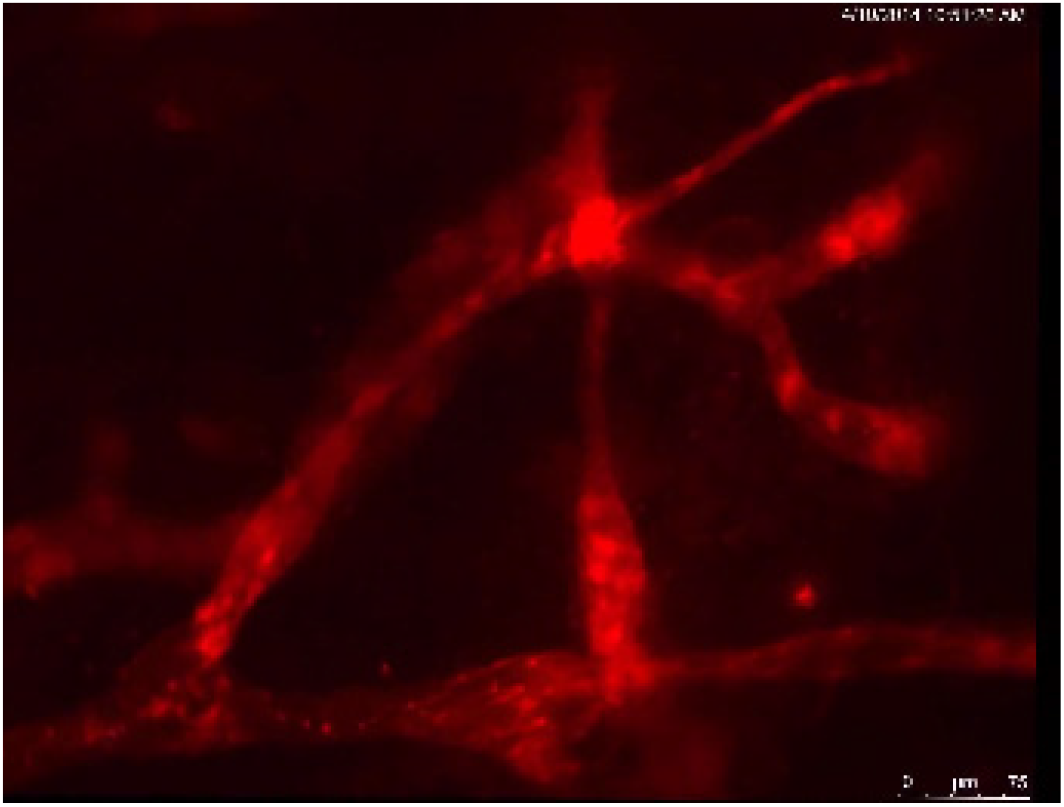
Lymphatic endothelium died off within two hours after photodynamic therapy. The video shows events from Figure 7A. Morphology of major collectors was identified with intravital staining for basement membrane (collagen IV) and co-expression of ProxTomato protein. For clarity, collagen IV staining is not shown on the video. As images were taken every one minute, Tomato-positive cells death appears as sudden ‘switching off’ (begins at the left side of the horizontal lymphatic) or gradual dimming (right side of the horizontal lymphatic). Cells’ budding (the indicator of apoptosis) was observed on endothelium treated with photodynamic therapy (PDT), while most cells die by necrosis. Cells dying by apoptosis do not lose their intracellular fluorescence intensity, and instead, their cytoplasmic proteins are compartmentalized into apoptotic bodies dispersing by Brownian movements (compare with Video 3). The imaging began 10 minutes after PDT. First cells started dying 50 minutes after PDT, and most cells within lymphatics were dead 2 hours later. The real-life duration of the video is 5 hours (timer jamming during video play are due to buggy Leica software export module). The video fps is 20 fps.

### Infiltration of DCs and T-cells within lymphatics persist throughout the regeneration process

Cessation of drainage after lymphatic decellularization is directly associated with the formation of intralymphatic clots of fibrin inside collectors but also massive infiltration of leukocytes **(Fig. 5E)**. Here we asked if the latter phenomena could participate in the occlusion of lymphatics. In addition, we wanted to know what cell types are represented in the immunological clusters that form within lymphatics.

In contrast to fast infiltration of leukocytes into ghost lymphatics after intradermal injection of directly toxic compounds, Triton or ferric chloride, milder decellularization of collectors with PDT resulted in a 24-hour lag phase and during that time leucocyte presence build-up only around fibrin occluded lymphatics **(Fig. 6A)**. Surprisingly, labeling of muscle cells smooth muscle actin with mouse monoclonal antibody revealed that intralymphatic clots could be directly stained with detection anti-mouse IgG antibody indicating that mouse IgGs derived from blood plasma are also trapped within lymph clots. Also, anti-mouse IgG staining also revealed that only a few IgG-expressing cells were present in the tissue of PDT-treated lymphatics, which was confirmed with anti B220 staining (not shown). The delayed infiltration of collectors is probably due to the preservation of afferent initial lymphatics and injection site, which in the case of Triton or ferric chloride injection, connect with collectors as one necrotic zone **(Fig. 5)**. This also indicated that it is fibrin clotting and not leukocyte accumulation within decellularized lymphatic directly followed endothelial cell death and was accurately correlated with lymphatic occlusion. Interestingly, the antigen-presenting cells **(Fig. 8B-C**) and T cells consist of the vast majority of cells that infiltrated ghost vessels while Gr1-positive neutrophils or monocytes and F4/80-positive macrophages and monocytes only populated ghost vessels-surrounding dermis (**Fig. 8C**). Antigen-presenting cells that populated ghost vessels expressed CD11c **(Supplementary Fig. 5)**, the marker of dendritic cells(Randolph,& al,2005).

**Figure 8.**
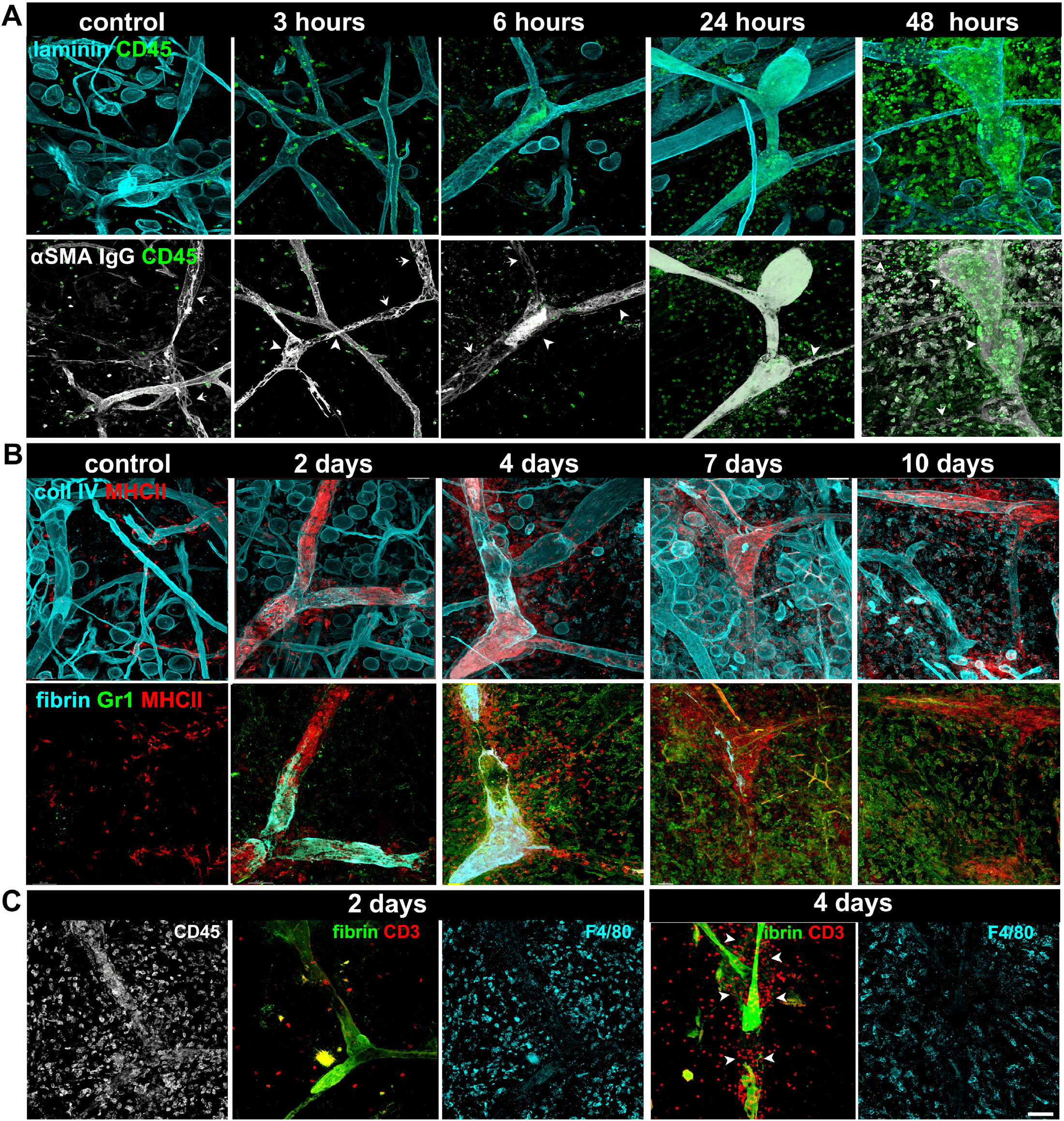
Massive infiltration of antigen presenting cells and T-cells into ghost vessels follows intravascular lymph clotting. **A**. Time course of leukocyte (CD45^+^ cell) infiltration into ghost vessels (as visualized by staining of laminin of basement membrane). Leukocyte accumulation within ghost vessels cannot be distinguished from that of control lymphatics for the first 24 hours after vessel decellularization, although leukocyte build-up in peri-lymphatic tissue is already apparent at 6 hours after PDT. A dramatic change in leukocyte accumulation within lymphatics occurs between 24 and 48 hours after PDT-dependent decellularization - lymphatics become densely filled with leukocytes making this leukocyte manifestation an identifier for decellularized vessels. Arrowheads point to intralymphatic protein clusters visualized with anti-mouse IgG antibody used to reveal mouse anti-αSMA staining. Arrows point to leukocytes clusters within ghost vessel. **B**. Dense leukocyte accumulation within lymphatic vessels persists for the whole regeneration time of lymphatic, observed up to 10 days. The majority of cells within lymphatics are MHC class II-positive antigen presenting cells. While Gr1-positive neutrophils and monocytes remained in the surrounding dermis. **C. 2 days** after decellularization. Single imaging field showing F4/80^+^ neutrophils and macrophages infiltrating dermis-surrounding collectors. However, they almost exclusively remain in the tissue and do not invade ghost vessels. **4 days** after decellularization. Single imaging field is showing that CD3ε+-T cells are the other abundant cell type that swarms ghost vessels by 4 days after decellularization. Arrowheads point to the border of a lymphatic collector marked by a residual sheet of fibrin, which delineates characteristic uneven diameter along the length morphology of lymphatic collecting vessels. Scale bar: 50μm.

**Video 2.**
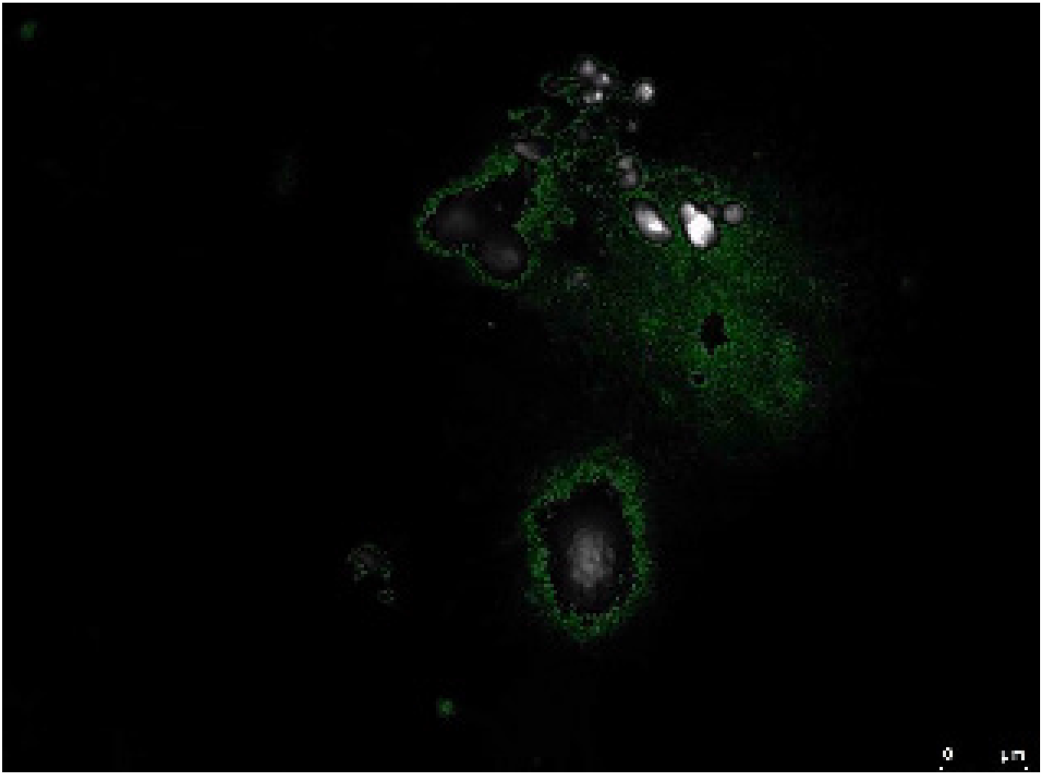
Concurrent necrosis and apoptosis of B16-F10 cells grown for one week in the mouse ear. Single cell divided but the two daughter cells died by apoptosis (top cell breaks into small apoptotic bodies which intensity is not different to the mother cell) and necrosis (bottom cell “explode,” the GFP fluorescence diffuse out in 1-2 frames of the 15 fps movie). The color gradient, from high-contrast white to green was chosen for the clarity. The real-life duration of the video is 2 hours. The video frame rate is 15 fps.

### Lymphatic endothelium is not required for fluid drainage and cell trafficking

Decellularization of collectors leaves an intact tube of basement membrane that is initially filled with impermeable fibrin clots to be later replaced by the basement-membrane-like matrix (**Fig. 6**). Since these ghost vessels were capable for at least short time drainage after decellularization, we asked whether lymphatic endothelium is at all needed for fluid drainage and traffic leukocytes to the lymph nodes. To do that, we tried to block lymph clotting by intradermal injection of heparin. This procedure was successful in 25% cases, as determined by the absence of fluorescence fibrin deposits (**Fig. 9B, live**). Without intravascular lymph clots decellularized lymphatics were able to drain dextran, however, their drainage was dysfunctional as they leaked collected fluid along their path. Nonetheless, even though not efficiently, lymphatics in the absence of endothelial lining can deliver macromolecules (e.g., antigens) to the draining lymph node. Furthermore, in the absence of endothelial lining and intralymphatic clots, remote segments of ghost vessels could be filled with erythrocytes, large particles that in contrast to fluid did not leak outside the basement membrane tube even 48 hours after injection (**Fig. 9B, fixed**). This indicates that endothelium is not necessary for cell passage to the draining lymph node. Interestingly, once ghost vessels were protected from the occlusion for 24 hours after the PDT, they stayed patent. Their draining state could be verified with lymphangiography even 12 days after PDT when all patent vessels drained dextran, but only collectors that recover endothelial lining were not permeable to the fluorescent marker already few minutes after the injection **(Fig. 9C**). Most importantly, decellularized collectors whenever they remained patent and drained fluorescent dextran were also capable of trafficking leukocytes, most importantly dendritic cells, to the draining lymph node (**Fig. 9D**), proving that presence of endothelial lining is not necessary for this process. At the same time, these results prove that occlusion of collectors with clotted lymph is sufficient to completely block the drainage of macromolecules (e.g., antigens) and trafficking of leukocytes from the tissue to the draining lymph node. Additionally, skin edema that develops one day after PDT was resolved on day 5 before any functional regeneration of lymphatics took place **(Supplementary Fig. 6)**. This is however in agreement with early observations, wherein the absence of persistent inflammation lymphatic occlusion alone does not produce lymphedema(Drinker,& al,1933).

**Figure 9.**
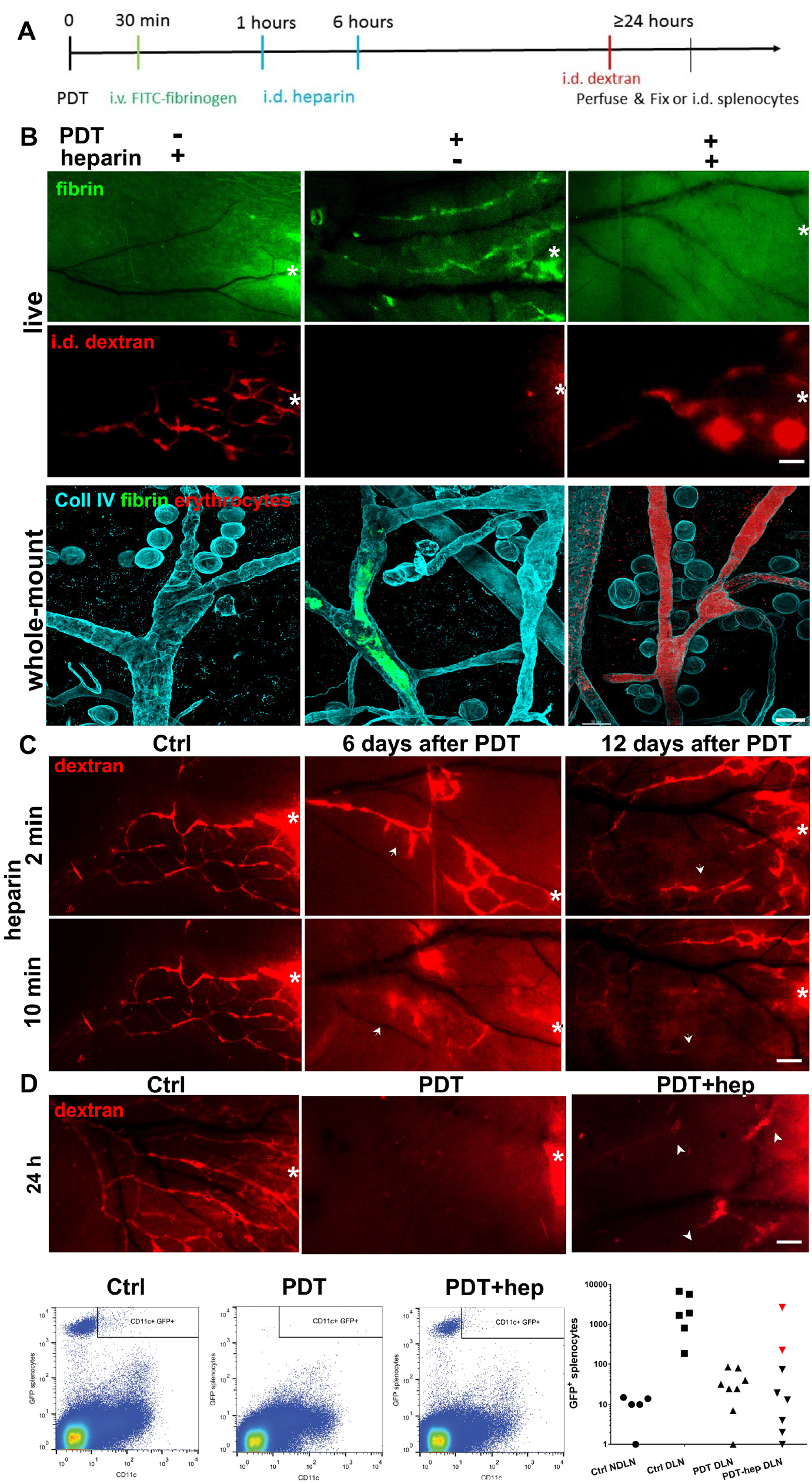
Heparin sustains lymphatic drainage in the deendothelized vessels. With inhibited clotting, decellularized lymphatic ghost vessels drain antigens and immune cells to the lymph node. **A**. Schematics of experimental design. Intradermal (i.d.) injection of Visudyne® mixed with 0.5 IU of heparin or saline is followed by collector imaging (verteporfin lymphangiography). Subsequently, the dorsal ear is irradiated with a 692nm laser (photodynamic therapy PDT). Thirty min later fluorescently labeled fibrinogen was injected intravenously (i.v.). Within the hour, and 6 hours after PDT, the same location in the ear skin was re-injected i.d. with 0.5 IU of heparin in 2.5 μl saline or saline alone. Eighteen hours later, fibrin deposits in the ear were imaged. Before fix-perfusion or at indicated time point functionality of ear lymphatics were assayed with fluorescent lymphangiography using intradermal dextran injection. **B**. Live imaging (live), **Top**: Heparin administration prevented intralymphatic clot formation along decellularized lymphatics. **Bottom**, lymphangiography: In the absence of intralymphatic clots, the decellularized lymphatics were able to drain fluorescent dextran. However, ghost vessels were permeable and leaked the tracer into the dermis. Whole-mount imaging (fixed). Decellularized lymphatics can take up large particles, demonstrated here with autofluorescent erythrocytes. By repeating injections of heparin, erythrocytes from non-coagulated blood enter the non-occluded, decellularized ghost vessels. **C**. Administration of two consecutive doses of heparin on the day of lymphatic decellularization was sufficient to maintain ghost vessel patency and drainage as seen at 2, and 10 minutes after i.d. dextran injection. Their leaky state persisted for 12 days. Arrows point to leaky lymphatic vessels. **D. Top**: Lymphatic function was assayed with fluorescence lymphangiography before injection of EGFP-splenocytes into the ear skin. Functional, PDT-decellularized lymphatic collectors remain patent with concomitant heparin injections but leaky. **Bottom**: Representative flow cytometry plots of EGFP splenocytes in the draining lymph nodes (DLN) and non-draining lymph nodes (NDLN) in control, and in DLN in PDT, PDT-heparin groups 24 hours after i.d. injection. Summary plot of flow cytometry EGFP+ cell counts. The red triangles in PDT-heparin group correspond to ears where heparin injection preserved dextran drainage as determined with fluorescent lymphangiography done prior splenocytes injection (**a**). Intensities of Gr1 and F4/80 were normalized to signal level of interstitial cells positive for these receptors. Scale bar: B-live; C, D-200 μm, B-whole-mount-50μm

**Video 3.**
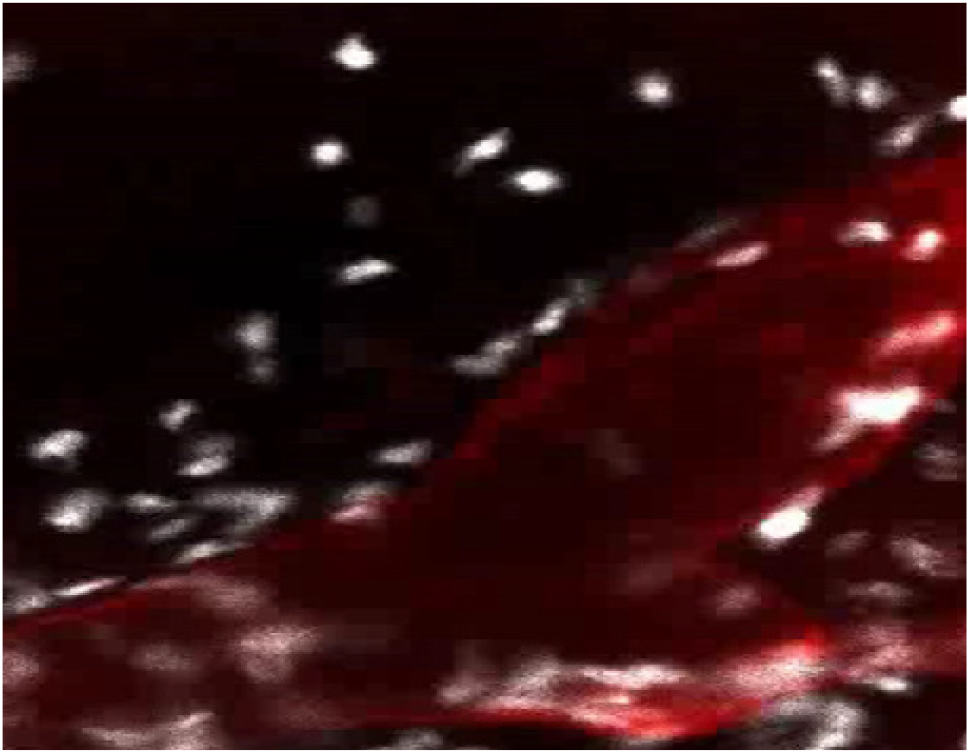
Necrotic death of lymphatic endothelium within the valve 1.5 hours after PDT. Propidium iodide, a membrane impermeable DNA-binding agent labels nuclei of necrotic cells. The real-life duration of the video is 8 minutes. The video plays at 4 fps.

### Lymphatic occlusion blocks immuno-recognition of allografts

Due to the potent and acute cytotoxic response and little effect of the immunosuppression on allograft tolerance, skin transplantations are confined to autografting(Benichou,& al,2011). However, cutaneous immunity has one significant vulnerability as its responsiveness depends on the functional lymphatic system(Lund,& al,2016) that connects the drained skin with secondary lymphoid organ(Lakkis,& al,2000). Allografts are accepted indefinitely when transplanted to a location that could not drain to the lymph nodes because of induction of immune ignorance rather than tolerance(Lakkis,& al,2000, Yamagami,& al,2001). Since we understood the occlusion mechanism and the kinetics of lymphatic regeneration, next, we planned to test the efficacy of the temporal lymphatic blockage on the prolongation of subcutaneous allograft acceptance. To do that we established a highly reproducible and sensitive subcutaneous heart transplant model(Fulmer,& al,1963). In contrast to skin-over-skin transplant assays, transplantation of self-contracting cardiac tissue provides a natural rejection parameter that can be visually assayed daily. Additionally, a pocket formed between cartilage and dorsal skin completely protected the implant from drying and infection and provided blood supply from the uninjured dorsal dermis. This assured a high level of the initial vascularization of the transplants (over 95%, Fig. 10A), which is the most critical parameter in avoidance of artifacts like considering necrotic rejection as an immunological rejection (McFarland,& al,2009). Transplanted under dorsal ear skin halves of the newborn mice’s hearts were vascularized within 5 days after implantation and started contraction that was monitored through the skin (**Video 4**). Control isograft or isograft transplanted under the PDT-treated ear skin were accepted until the end of the experiment (**Fig 10B**). In contrast allografts, the median rejection time was 8 days (**Fig. 10B**). The average graft survival was extended to 12 days when allografted hearts were implanted under PDT-treated skin. In addition, the rejection kinetics in the control and PDT groups were different. While the control tissues were rejected across the 5 day period, the rejection in PDT group is almost binominal and occurred within one day. This could be explained by the high level of lymphatic regeneration homogeneity (**Fig. 10C**) but also the fact that ghost vessels before the regeneration of lymphatics become tertiary lymphoid organs(Ruddle,2014), with an accumulation of antigen-presenting cells ready to be delivered to the draining lymph node (**Fig. 8**). Conceivably, as the time of vessels regeneration was delayed with anti-VEGFR3 therapy (**Fig. 10C**), the time of allograft acceptance was almost doubled from the original 6 to 15 days. Interestingly, this value is better than presented with the same assay results on the effect of commercial cyclosporine (Sandimmune) treatment on allograft rejection that occurred after 9 days in the control and 13days cyclosporine-treated group(Gorecki,& al,1991).

**Figure 10.**
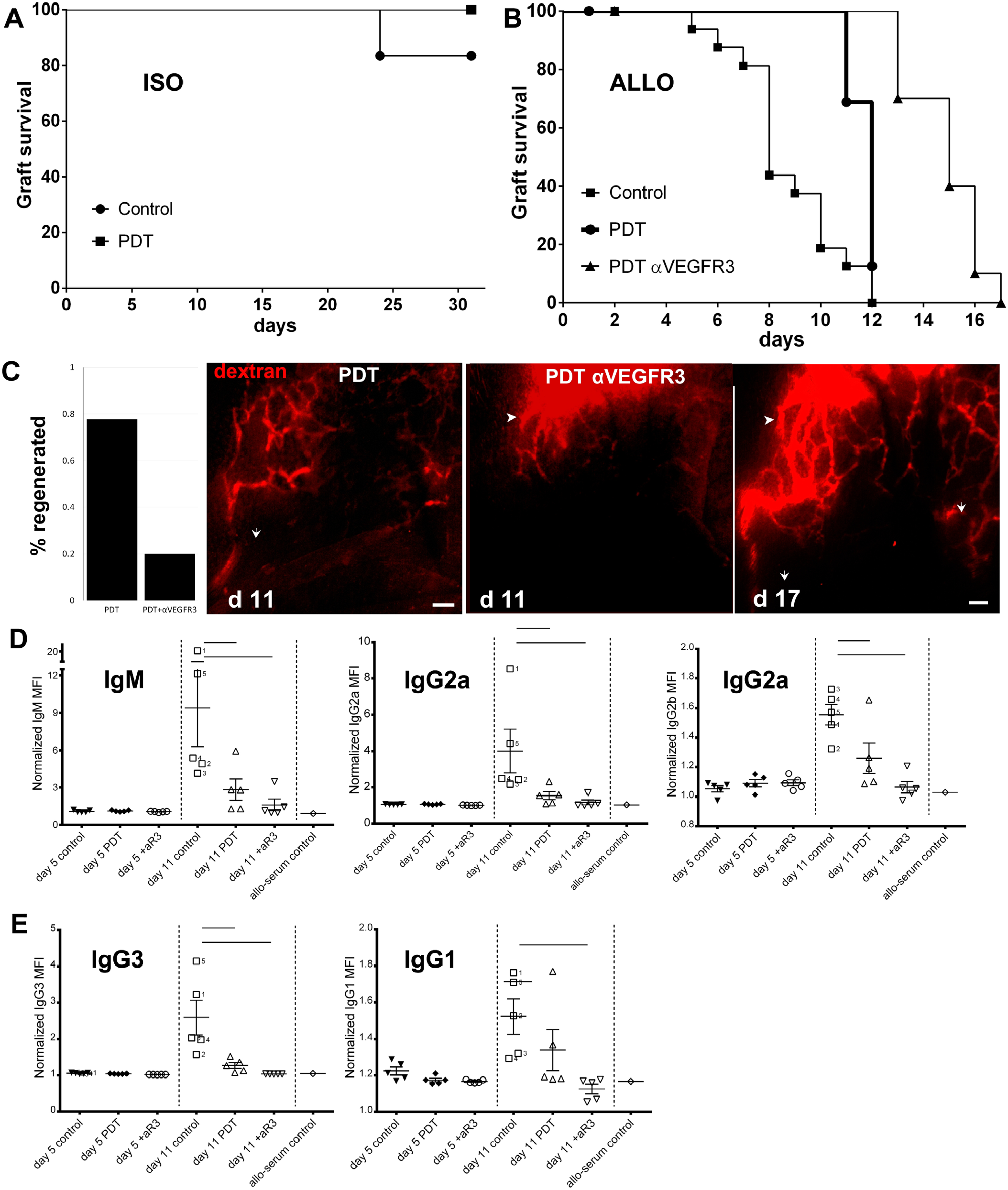
Lymphatic occlusion hides antigens and prevent rejection of skin allograft. Murine embryonic contractile cardiac tissue was transplanted under ear dermis of the fully immunocompetent mouse with or without lymphatic-specific photodynamic therapy (PDT) and observed for functional muscle contraction, plotted as % graft acceptance rate vs. number of days of graft survival (**A, B**). **A**. Isografts from control group (n=6, 5 censored) and PDT group (n=6, 6 censored) continued contracting until the planned termination of the experiment (31 days) where lymphatic collector decellularization with PDT had no effect on isograft acceptance. (Mantel-Cox log-rank test (p<0.32). All implanted hear fragments resume contraction after implantation. **B**. The median survival time of allografts transplanted under control mouse dermis was 8 days (n=17 mice, 16 rejected, 1 censored). When allografts were implanted under immuno-priviliged ear dermis with PDT-occluded lymphatics, the kinetics of allograft rejection was uniform, and graft median acceptance was prolonged to 12 days (n=18 mice, 14 rejected, 4 censored). This median acceptance time of the allogeneic transplant grafted under PDT-decellularized lymphatics was prolonged to 15 days (Mantel-Cox log-rank test (p<0.0001) with adjuvant anti-lymphangiogenic therapy with intraperitoneal administration of anti-VEGFR3 antibody (PDT αVEGFR3 group, n=11 mice, 10 rejected, 1 censored). All censored cases in allograft groups were due to failure to resume contraction during the first 5 days after the implantation. **C. Lymphangiography**, quantification. 7 out of 9 ears harboring subcutaneous implant resumed lymphatic drainage 11 days after treatment in PDT group. In contrast, mice receiving adjuvant αVEGFR3 antibodies after PDT-induced lymphatic decellularization had only 2 out of 10 ears that resumed drainage (p=0.023, Fisher exact test). Representative lymphangiographic images showing drainage at day 11 after PDT in PDT and PDT+αVEGFR3 ears and the drainage in the same PDT+αVEGFR3 ear on day 17. Arrows show the direction of regenerated drainage. Arrowheads on day 11 and day 17 point to the same lymphatic vessels. Scale bar: 250μm. **D**. Titers of complement activating (antibody-dependent cytotoxicity, ADCC) and phagocytosis-activating alloantibodies (cell-dependent cytotoxicity, CDC) of all classes in the sera of mice in control, PDT and PDT αVEGFR3 groups at day 5 (before the formation of antigen-specific antibody response) and day 11 after PDT. The titer of allo serum control is shown as a point of reference. Bars indicate the difference between groups tested with ANOVA and Holm-Sidak post-priori test at p<0.05.

**Video 4.**
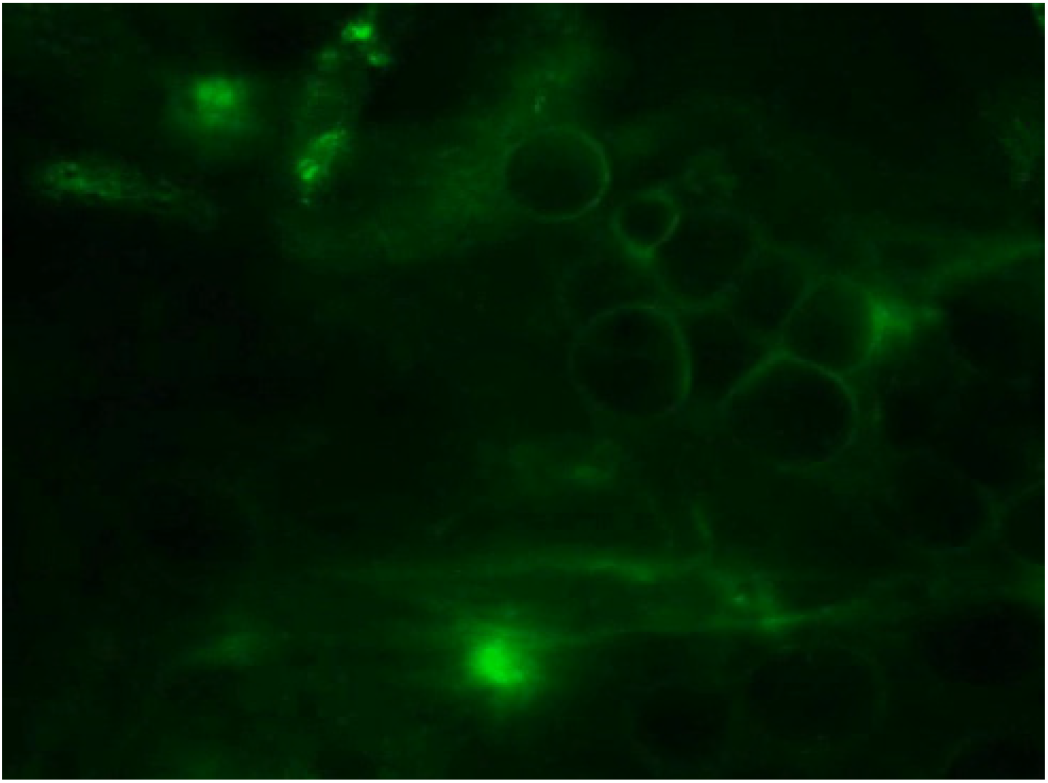
Fibrin deposition on the wall of collecting lymphatic 2 hours after PDT. Imaging that started 2 hours after PDT and 1.5 hours after intravenous injection of FITC-fibrinogen was continued for 2 hours with images taken every minute. Microinjuries that are the effect of surgical exposition of dorsal dermis lead cased generalized leakage of plasma from vessels leading to an increase in fibrinogen fluorescence outside blood vessels. Immediately after PDT, mouse ear dorsal skin was exposed, and dermis was stained with collagen IV (not shown) to identify draining lymphatic vessels based on their characteristic morphology. The real-life duration of the video is 2 hours. The video plays at 15 fps.

**Video 5.**
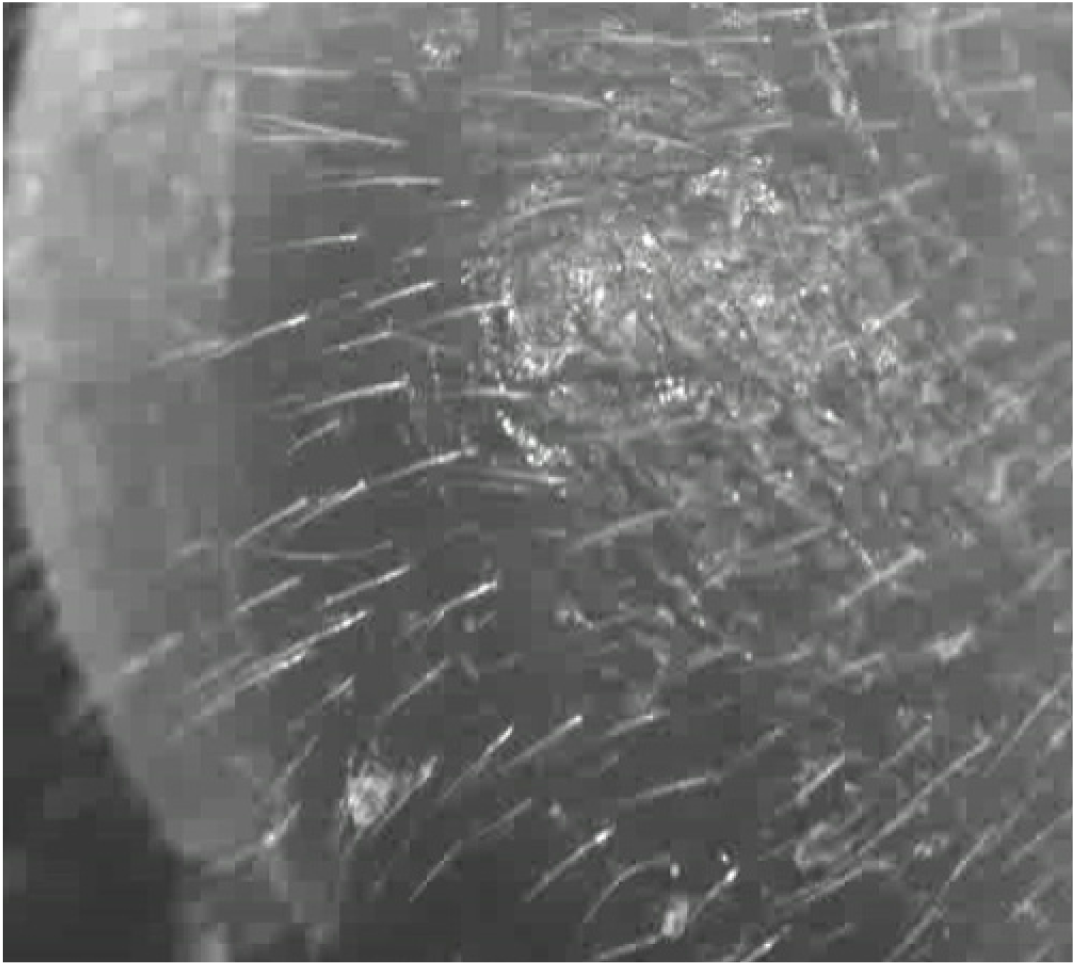
Cardiac muscle regular contraction after transplantation under mouse ear skin. A newborn mouse heart fragment transplanted under the skin of the mouse ear started beating after 3-4 days and continues as long the implant-specific immune response is prevented. Images were taken every 0.2 seconds for 10 seconds. The video plays at 15 fps (speed up 2x).

Alloantigen-specific antibody that develops after the tissue implantation is generally considered not relevant for an acute cytotoxic response. Yet, their level and subtype constitute significant parameters in probing immune ignorance as to the production of allospecific antibodies, and antibody class switching reflects the availability of alloantigens in the lymph node(Colvin,& al,2005). The seven-fold relative increase of the earliest released IgM alloantibody was the highest among all tested isotypes (Fig. 10C) indicating that anti-lymphatic (PDT) therapy reduced the increase of allospecific antibody to 1.4 fold (2.4-times), with a further reduction to the basal level (0.4 fold increase) was measured when anti-lymphatic therapy (PDT) was followed by lymphatic regeneration delay with αVEGFR3 therapy. Even though the level of antibodies in PDT and PDT-αVEGFR3-treated groups were not different from the respective basal levels, the patterns of antibodies decline were reproduced for every antibody subtype (five curves not different, p>0.05) in PDT-only and PDT+αVEGFR3 groups. This means that rather than the development of immune tolerance, the immune reaction was only delayed and that alloantigens that remain outside the reach of the lymph node did not lead to T-cell tolerance. Instead, these alloantigens are ‘ignored,’ and the alloreactive T cells remain functionally intact and can be engaged when tissue clearance is restored.

## Discussion

Here we characterize the anti-clotting properties of the lymphatic collecting vessels and their application in the on-demand cessation of antigen drainage and trafficking of the antigen-presenting cell. This leads to the creation of subcutaneous immune-privileged zones where allografts can be protected from immune recognition without immunosuppression. We show that fluid drainage and immune cell trafficking entirely depend on the lymphatic vessel ability to sustain lymph liquidity.

### The physiological role of lymphatics hemostasis

Regardless of the method of lymph clotting induction, intravascular lymph clots interfered with essential lymphatic functions, fluid, and macromolecule evacuation from the drained area of the skin. Intralymphatic clots were detected by copolymerization of fluorescently labeled fibrinogen. Because FITC-fibrinogen was injected intravenously, first it had to leak with other plasma proteins to the tissue (at the site of prior intradermal injections). Later these proteins were collected by lymphatics and drained as naturally formed lymph. Therefore the fluorescent labeling of intralymphatic clots reflected the natural process of lymph coagulation.

In contrast to the selective induction of pro-hemostatic TF expression that resulted in discrete and sparse lymph clots deposition within collecting lymphatics (Fig. 1-2), inhibition of constitutively expressed thrombomodulin resulted in widespread lymph clotting inside efferent vessels (**Fig. 3B and 4B, C**). These lymph clots formed within intact lymphatic intima were cleared within subsequent 16 hours, therefore, their physiological or therapeutic relevance is limited to extemporary, e.g., inhibition of bacterial spread in early acute inflammation, as previously reported(Menkin,1931, Menkin,1931, Menkin,1931). Similar to thrombomodulin inhibition, decellularization of lymphatic intima lead to massive and continuous lymphatic occlusion independent on LEC expression profile or their location within the collector, leaving ‘ghost vessels’ composed of acellular basement membrane filled with lymph clots (**Fig. 5-6**). Experiments with an injection of necrotizing compounds, i.e. Triton X-100 and FeCl_3_ (**Fig. 5**) aimed to confirm previous observations where intradermally administrated turpentine or FeCl_3_(Menkin,1931, Menkin,1931) lead to cessation of lymphatic drainage by intralymphatic lymph clotting (referred as ‘thrombosis’, experiments reviewed by Drinker and Field(1933)).

Both, Triton X-100 and FeCl_3_ exerted toxicity at the site of injection but also completely blocked the skin drainage and induced lymph clotting in the lymphatics located remotely to the site of intradermal injection. The same effect of lymphatic occlusion but without adverse skin, necrosis was achieved with PDT-induced lymphatic decellularization (Fig. 6-10). Due to the high specificity of anti-lymphatic treatment and the long-lasting (9 days) drainage occlusion, we used this method to study the effects of intravascular lymph clotting on lymphatic physiology and local immune responses.

Despite the type of lymphatic treatment, lymph clotting was an endpoint that preceded interference or complete blockage of fluid drainage. Even though leukocytes involvement in fibrin polymerization and maturation within lymphatics could not be excluded; the initial lymph coagulation occurred before leukocytes infiltration (Fig.2 and 8). Therefore, in one way or another lymph clotting was dependent on lymphatic endothelium that either expressed TF after stimulation lost anti-hemostatic properties after thrombomodulin blocking or was destroyed by detergent lysis (Triton-X-100), exposure to free-radicals generated by FeCl_3_ or free-radicals generated by light-activated verteporfin (i.e., lymphatic-specific PDT). However, the duration of lymphatic occlusion varies significantly between the metabolic and noxious methods of lymph clotting, from less than 22 hours with preserved intact endothelium to 9 days(Kilarski,& al,2014), when collectors were decellularized. Initial lymph clotting could be activated by dying cells releasing pro-thrombotic negatively charged nucleic acids and polyphosphates that can activate factor XII of the contact-dependent (intrinsic) coagulation pathway(Esmon,2010). The intrinsic pathway can be also continuously activated by the constant exposure of lymph to the unmasked basement membrane collagens ((van der Meijden,& al,2009) and **Fig 6C-D**), which in the absence of endothelial membrane-bound antithrombotic thrombomodulin((Maruyama,& al,1985) and **Fig. 1A**) can lead to recurrent activation of lymph coagulation even when the initial clot is cleaved 4-5 days after PDT-driven decellularization of lymphatics (**Fig. 6F**). The stability of intralymphatic clots is additionally enhanced by the loss of the endothelium that is a source of tissue plasminogen activators(Laschinger,& al,1990), which production is enhanced after pro-thrombotic stimulation of endothelium (**Fig. 3D**). Therefore, restoration of lymphatic intima should be the critical factor in the stable preservation of collector patency. This conclusion agrees with our subsequent finding that lymph clotting but not the loss of endothelial lining was a direct cause for lymphatic occlusion to fluid and cells. This is indicated by a delayed block of drainage that occurred hours after endothelial death (Fig. 7), and most notably by sustained fluid drainage and cell trafficking through PDT-decellularized collectors when fibrin deposition was inhibited with heparin (Fig. 9). Even though decellularized collecting lymphatics were leaky to drained dextran, these experiments indicated that endothelial lining is not indispensable for the most affirmed functions of lymphatics, the fluid drainage, and cell trafficking and instead lymphatic endothelium is primarily needed to support the collector’s patency.

The most striking conclusions from our experiments pointed to the role of lymphatics in quenching the continuously self-activating clotting system in homeostasis and terminating the thrombin-dependent amplification phase of the clotting, limiting it to the local inflammatory sits. In this sense, lymphatic function as the ideal the systemic spread of clots from.

### Lymphatic anti-clotting mechanism in prevention of allograft rejection

#### Organ rejection

This lymphatic function might explain the puzzling presence of lymphatics within organs like the kidney and the heart, where predominantly blood circulation maintains tissue-blood fluid balance(Guytron,& al,2016). During transplantation, lymphatic collectors in these organs are either ligated or left untied, with no apparent outcome on the organ health(Kilarski,2018). However, lymphatic quenching of fibrin formation could preserve the highly structuralized organizations of these tissues from imbalanced clotting and subsequent fibrosis, potentially contributing to the organ rejection. Increasing the resilience of intralymphatic clots could also extend non-anastomosed (e.g., endocrin) subcutaneous allografts that are masked and ignored by the immune system(Kilarski,2018).

#### Subcutaneous tansplantation of endocrine tissues

The lack of conventional lymphatic drainage is a characteristic feature shared between passive immune ignorance(Lakkis,& al,2000) and immune privilege at sites like the placenta, cheek pouch, cornea, or central nervous system (Medawar,1948, Simpson,2006, Aspelund,& al,2015). Furthermore, it was shown that surgical separation of draining lymphatics blinds the draining lymph nodes to antigens residing within the blocked area and prevents the development of adaptive immune response against allograft at least for the time alternative lymphatic routes developed (Barker,& al,1968, Lambert,& al,1965). Due to extensive and traumatic surgery, these experiments lack translational potential. Instead, they provided a proof of concept where elimination of antigen drainage path leads to the formation of an immune privilege zone where foreign antigens remain ignored by adaptive immunity. Long-lasting occlusion of lymphatic collector that follows specific eradication of their intima allowed us to mimic the effect of mechanical laceration of the skin lymphatics, proving the therapeutic applicability of the anti-hemostatic properties of lymphatic endothelium. Instead of an ectopic skin transplant, where rejection is defined by subjective parameters(Joffre,& al,2008), we used a sensitive cardiac graft where the immune rejection is manifested by the loss of spontaneous contractile function(Fulmer,& al,1963). Indeed, the rejection of allografts but also the development of allospecific antibodies was delayed after PDT-dependent decellularization of collectors and correlated with drainage recovery from the skin surrounding the graft (Fig. 10). Importantly, the time of graft acceptance was doubled as the immune response was delayed when lymphatic regeneration was inhibited with VEGFR3 blocking antibodies. The possibility to further prolong allograft hiding from the immune recognition with anti-lymphangiogenic therapy indicates that there are still opportunities for further enhancements in the adjuvant therapies, e.g., by reinforcing clotted lymph with matrix-bound protease inhibitors(Lorentz,& al,2011). Interestingly, in the two combined relatively non-invasive treatments, lymphatic decellularization with PDT and subsequent inhibition of lymphatic regeneration with anti-VEGFR3 antibody, had a stronger effect on rejection inhibition (delay by 47%) than immunosuppressant cyclosporine in a form of clinically-approved Sandimmun® (delay by 30%), when tested in cardiac allograft model(Gorecki,& al,1991). However, it is important to appreciate the difference between these treatments, while lymphatic occlusion leads to immunological ignorance by hiding tissue from immune recognition in draining lymphatics, cyclosporine suppresses already developing an immune response. Accessibility of the dermal lymphatics and low-risk subcutaneous surgery might permit therapeutic prosthetic transplantation of small hormone-producing various tissue fragments or cells, like Langerhans islets (Pepper,& al,2015).

### Potential role of lymphatic anti-clotting in pathology

#### Lymphedema and inflammation

We described the overlooked function of lymphatic vessels that might have significant consequences in various pathologies and therapies that until now could not be adequately appreciated. In homeostasis anti-hemostatic mechanisms helps sustain lymph as a liquid but during the acute inflammation, the protein-rich exudate that leaves blood vessels does not enter lymphatics, helping to circumscribe the infection from draining to the efferent vessels and eventually bloodstream, reviewed by Menkin (1931) and Drinker (1933). This phenomenon of ‘fixation’ of bacteria, protein, and particles to the site of early inflammation was attributed to the mechanical obstruction of lymphatics with fibrin leaking from inflamed or injured blood vessels. Unfortunately, as it could explain, e.g. why acute lymphangitis, an infection of lymphatic collectors is predominantly caused by plasmin-activating positive group A streptococci bacteria(Ferri,2014) the research on in situ lymph clots was not further studied(Menkin,1960). Furthermore, the ability of dysfunction of lymphatics to sustain non-clotted lymph might contribute to their pathologies with idiopathic or confound origin. Surprisingly, even in the case of thoroughly studied lymphedema, causative factors leading to the condition are not known as lymphatic occlusion is a necessary but still, insufficient factor to induce sustained lymphedema in animal models(Drinker,& al,1933, Rutkowski,& al,2006) but also in post-mastectomy patients(Rockson,2021). Also, lymph clots within lymphatics around the granuloma that form after worm death during the pathogenesis of filarial lymphedema(Case,& al,1991) intuitively carry a notion that lymph clotting and inflammation might be important priming factors of the disease. Filaria-induced lymphedema develops after repeated instances of worm deaths and bacterial lymphangitis that additionally lead to lymphatic endothelium injury and fibrotic remodeling of lymphatic collectors(Chakraborty,& al,2013). Along these lines, our results indicated that inflammatory factors-stimulated lymphatic occlusion is transient and quickly resolved within lymphatic vessels, while injury to lymphatic endothelial lining is sufficient for prolonging occlusion of the collector. But even after PDT-induced lymphatic decellularization, ghost vessel eventually regenerated instead of becoming fibrotic(Kilarski,& al,2014). PDT-decellularization is a specific and sterile anti-lymphatic treatment while trauma combined with infectious inflammation might produce long-lasting intralymphatic clot remodeling and permanent vessels occlusion with collagenous fibrosis, elements of which we could observe in lymphatics regenerating after PDT (Fig 6F). Therefore, anti-hemostatic properties of lymphatics should be the primary parameter that requires consideration before designing lymphatic implants, e.g., artificial collectors lympho-venous shunts or lymphatic collateral neovessels(Patel,& al,2015) as they must anchor functional anti-hemostatic mechanism in order to canalize the clottable fluids drained from edematous tissues. Because the anti-hemostatic property of lymphatic received little attention, it is possible that lymph clotting and subsequent fibrosis is an underestimated complications unwittingly hampering lymphedema treatment. For example, unpredictable clotting and fibrinolysis cycles could explain the fact that most silicon tubes implanted in lymphedema patients did not drain the fluid along their lumen even though it did not undergo fibrosis and remained patent(Olszewski,& al,2015).

#### Skin infections and metastasis

The active regulation of lymphatic drainage might also affect infection spread and tumor metastasis. Entrapped tumors or associated cells might activate fibrinolytic mechanisms and use residual fibrin scaffolds to enhance their migration within the lymphatic system. This scenario is supported by experiments where fibrinogen depletion inhibits lymphatic metastasis but has no effect on the primary tumor growth (Palumbo,& al,2002). Contrarily, completely occluded lymphatics might stop the spread of metastatic cells, likely the mechanism of PDT treatment that blocked the metastasis spread via the peri-tumoral lymphatics (Tammela,& al,2011). Reinforcement of lymph clots by, e.g., local inhibition of fibrinolysis with fibrin-binding aprotinin(Lorentz,& al,2011) could prolong an inhibition of cell or pathogen spread(Menkin,1931).

#### COVID-19

Aberration in the lymphatic control of clotting and fibrinolysis might also provide clues on the post-infection COVID-19 pathologies that have a disastrous effect on remote but never infected organs(Loo,& al,2021). For instance, the premature but continuous release of lymph clots from small collectors in the lungs to the venous circulation could occlude the capillary system of distant organs, amplifying their pre-existing pathologies, leading to inflammation and, in the end, to organ failure. Notably, the prolonged discharge of micro clots from the thoracic duct could explain vascular damage in CNS and frequent mental abnormalities, even though the brain is rarely infected by COVID-19(Achar,& al,2020).

## Methods

### Animal experiments

Animal experiments were performed at the EPFL (Lausanne, Switzerland), the University of Chicago and the Medical University of Warsaw (tail edema experiments). Procedures performed on animals were in strict accordance with the Swiss Animal Protection Act, the ordinance on animal protection and the ordinance on animal experimentation. We confirm that our Institutional Animal Care and Use Committee (IACUC), named Commission de Surveillance de l’Etat de Vaud (Permit Number: 2646 and 2687), specifically approved this study or in accordance with protocols approved by the Institutional Animal Care and Use Committee at EPFL (permit number 72414). Procedures performed at the University of Chicago were performed according to rules and protocols at the University of Chicago were performed according to the guidelines approved by the Ethics Committee of the Medical University of Warsaw. BALB/c female or C57 mice were purchased from Jackson Laboratories or Charles River, maintained and bred in a specific pathogen-free barrier facility, and used between 6 to 10 weeks of age unless stated otherwise.

### Anesthesia and hair removal

Three days before each experiment, ears were treated with depilation cream (Veet® Hair Removal Cream, Slough, UK) for 10 s followed by thorough rinsing with water. For PDT, mice were anesthetized with 2.5 % isofluorane (Rothacher GmbH, Bern, Switzerland) and kept under 1.5 % isoflurane for the duration of the experiment. Mouse core temperature was maintained at 37 °C throughout the experiment (DC Temperature Control System, FHC Inc., Bowdoin, MA). All intradermal injection in the dorsal mouse ear (0.5 μl) was done using a microsyringe (Hamilton, Reno, NV, USA) equipped with 33 G needle.

### Lymphatic specific photodynamic therapy

The mouse head was epilated under isoflurane anesthesia 24 h before the procedure. Ears were treated with epilation cream (Veet) briefly for 10 s at the end of the head epilation process, and the mouse head was washed with plenty of water. The mouse was allowed to recover in a cage with food and water ad libitum. On the day of the experiment, a mouse was anesthetized with 2.5% isoflurane and kept under 1.5% of isoflurane for the duration of the experiment. 0.5 μl verteporfin in the form of liposomes (Visudyne®, Novartis Ophthalmics, Hettlingen, Switzerland) freshly reconstituted from the powder as previously described(Kilarski,& al,2014) was injected in the top of the ear with Hamilton syringe (Reno). Injection site of the ear was covered by aluminum foil and 1 min later, the ear was irradiated with non-thermal laser light (Biolitec) or 2W LED laser (Modulight) at 689 ± 2 nm delivered through an optical fiber containing a frontal light distributor with a lens (Medlight) with the irradiance of 50 mW/cm2, and a light dose of 100 J/cm2. Light doses were adjusted with neutral density filters and measured with a calibrated Field-Master GS power meter (Coherent).

### Decellularization of lymphatics with detergent and ferric (III) chloride

5% Triton-X-100 (TX100) and FeCl_3_ (Ferrum chloride) were dissolved in water and filtrated through the 0.22μm membrane. 0.5μl of TX100 or Ferrum chloride was injected in two adjacent spots at the top of the ear dorsal dermis and immediately after that FITC labeled fibrinogen was injected i.v.

### Fluorescent labeling of fibrin

40 ml 10 mg/ml fibrinogen (SIGMA) dissolved in in PBS was reacted with 1.6 ml of 4 mg/ml FITC (ThermoFisher) for 1 hour and fibrinogen was dialyzed 4 times to 4L each time of PBS. After that fluorescent fibrinogen was aliquoted and stored at -80°C until used. After thawing 200 μl of labeled fibrinogen was injected i.v. into mouse tail vain up to 15 to 30 minutes after treatment that was expected to induce intralymphatic coagulation (PDT, thrombin, cytokine injection).

### Thrombomodulin blocking and induction of TF expression on lymphatics

1 U/μl thrombin was reacted with 1 mM phenylmethylsulfonyl fluoride (PMSF, Sigma) for 2 hours and residual PMSF was removed on the Zeba™ desalting column (ThermoFisher Scientific). This treatment renders 85% of thrombin inactive as determined with the delayed fibrin clotting (from 45 seconds to 270 seconds).

Complete inactivation of thrombin was done by reacting of 50 μl of 1 U/μl thrombin with 1 mM p-amidinophenylmethylsulfonyl fluoride (p-APMSF, EMD Millipore Corp. (Laura,& al,1980)) μl of thrombin for 2 hours and residual PMSF was removed on the Zeba™ desalting column. 0.5μl of inactive thrombin or partially inactivated thrombin was injected intradermally alone or with 10ng/μl TNFα or 10νg/μl Il-1β in the ear top dorsal dermis.

### Splenocytes trafficking into lymphatics

Splenocytes from BALB/c mouse expressing eGFP under ubiquitin promotor were isolated from the mouse spleen by grounding the spleen through the strained with 70μm pores. Red blood cells were lysed with RBC lysing solution (150 mM NH_4_Cl and 10 mM NaHCO_3_) for 5 minutes, washed twice in DMEM and dissolved in Ringer’s buffer buffered with 25 mM HEPES, pH 7.5. Cells were spun down for the last time, loaded to the Hamilton syringe with the pipette tip. After mounting 33G needle, 2.5 μl slurry was injected into the same location of the ear where verteporfin and heparin or Ringer was injected on the previous day.

### Fluorescent lymphangiography

Two hours after PDT and every 2-3 days 1 μl of TRITC-dextran (10 μg/ml) was injected in the location of initial verteporfin injection. Draining lymphatics were imaged with stereomicroscope after the ear was placed between two glass slides. For intravital lymphangiography, ventral skin with underlying cartilage was separated and removed from dorsal, PDT-treated skin and primary rabbit, anti-collagen IV antibody, was applied for 20 minutes. After brief washing with Ringer’s buffer, secondary streptavidin-647 was applied for 5 minutes; tissue was washed and imaged using automated fluorescent stereomicroscope (M205 FA, Leica Microsystems) coupled to the sensitive color camera Leica DFC350 FX (Leica Microsystems). The following objectives have been used: Plan Apo 1x, NA 0.035; W.D. 60 or Plan Apo 2x, NA 0.07; W.D. 60 (Leica Microsystems).

### Whole-mount staining of a mouse ear

Mice were sacrificed with CO2 and exsanguinated by intracardiac perfusion with 20 ml Ringer’s buffer (supplemented with 10,000 IU/L heparin, 0.1% glucose, and 25 mM HEPES; pH 7.5, 330 mOsM) at a constant gravitational pressure of 120 mm Hg. Ringer’s buffer was then changed to osmolarity-corrected zinc fixative (Zn fix) solution [26] (4.5mM CaCl2, 52 mM ZnCl2, 32 mM Zn(CF3COO)2, 2mM Triss, 38 mM glycine; pH 6.5, 340 mOsm/L) and ears were cut and placed in ice-cold Zn fix with 1% Triton x-100 for at least 24 h. After that, the ventral part of the skin was removed together with cartilage and muscles, and dorsal dermis was washed in TBS (140 mM NaCl, 25 mM Triss, pH 7.5) for 6 h, blocked with 0.5% casein in TBS (blocking solution) for 2 h, and incubated with anti-VE-cadherin, anti-Lyve-1, and biotinylated anti-collagen type IV antibodies (10 μg/ml) in blocking buffer for 24 h. After washing in TBS with 0.1% Tween 20, tissue was incubated with the appropriate secondary antibody (10 μg/ml) for another 24 h, washed in TBS and dehydrated with 70% and 100% ethanol before it was cleared with 2:1 benzyl benzoate/benzyl alcohol solution supplemented with 25 mg/ml propyl gallate (refractive index 1.56). The tissue was mounted on a glass slide and imaged using HC PL APO 20x, NA 0.70 or HCX PL APO 63x, NA 1.40 lenses of a Leica SP5 confocal microscope (Leica). Stacks of images were analyzed with Imaris 7.4 (Bitplane AG).

### Subcutaneous graftin of cardiac muscle

PDT was performed on ears on day 0 and on the next day newborn (1-2 days after birth) mice hearts from allogenic or syngeneic mice were cut in half and incubated in DMEM media. The incision was done on the ventral side of the mouse ear and spatula was inserted through the cartilage to the end of the dorsal, separating cartilage from dorsal ear skin. Inside the forming chamber, halve ear was inserted, and the ventral incision was sealed with surgical glue. 50 μl of the blood was collected on day 5 and 10, and isolated serum was frozen for further analysis. Heartbeat in mouse ear was monitored daily after short isoflurane anesthesia. Until the beating stopped when mice were perfused, and their ears were zinc-fixed for further analysis.

### Detection of allospecific IgM and IgG in grafted mice

Thymocytes from the allogenic background were isolated from 4 weeks old mice at 10×10^6^/ml, and 50μl of cells were mixed with 5μl of mouse serum isolated from mice bearing transplants on days 5 and 10. After washing, cells were incubated with appropriate isotype antibody or anti-IgM antibody, re-suspended for FACS buffer for acquisition in Beckman Coulter CyAn ADP flow cytometer.

## Acknowledgments

The authors are grateful to Melody Swartz and Angelika Muchowicz for help in planning and performing the experiments, S. Hirosue for editing the manuscript and advice, V. Triacca, E. Guc, R. Mężyk-Kopeć, Jakub Gołąb and J. Kilarska for their technical assistance. The authors are grateful to the EPFL Bioimaging and Optics Platform and Histology Core facility for their technical assistance. Funding for this project was provided in part from the European Commission Framework Project 7 Angioscaff, and from the European Research Commission Advanced Grant LymphImmune, 323053.

## Author contribution

M. Wachowska performed and planned experiment. WW Kilarski conceived the idea, planned experiments, analyzed data and wrote the manuscript.

## Conflict of interest

Authors declare no conflict of interest.

---

**Supplementary Fig. 1.**
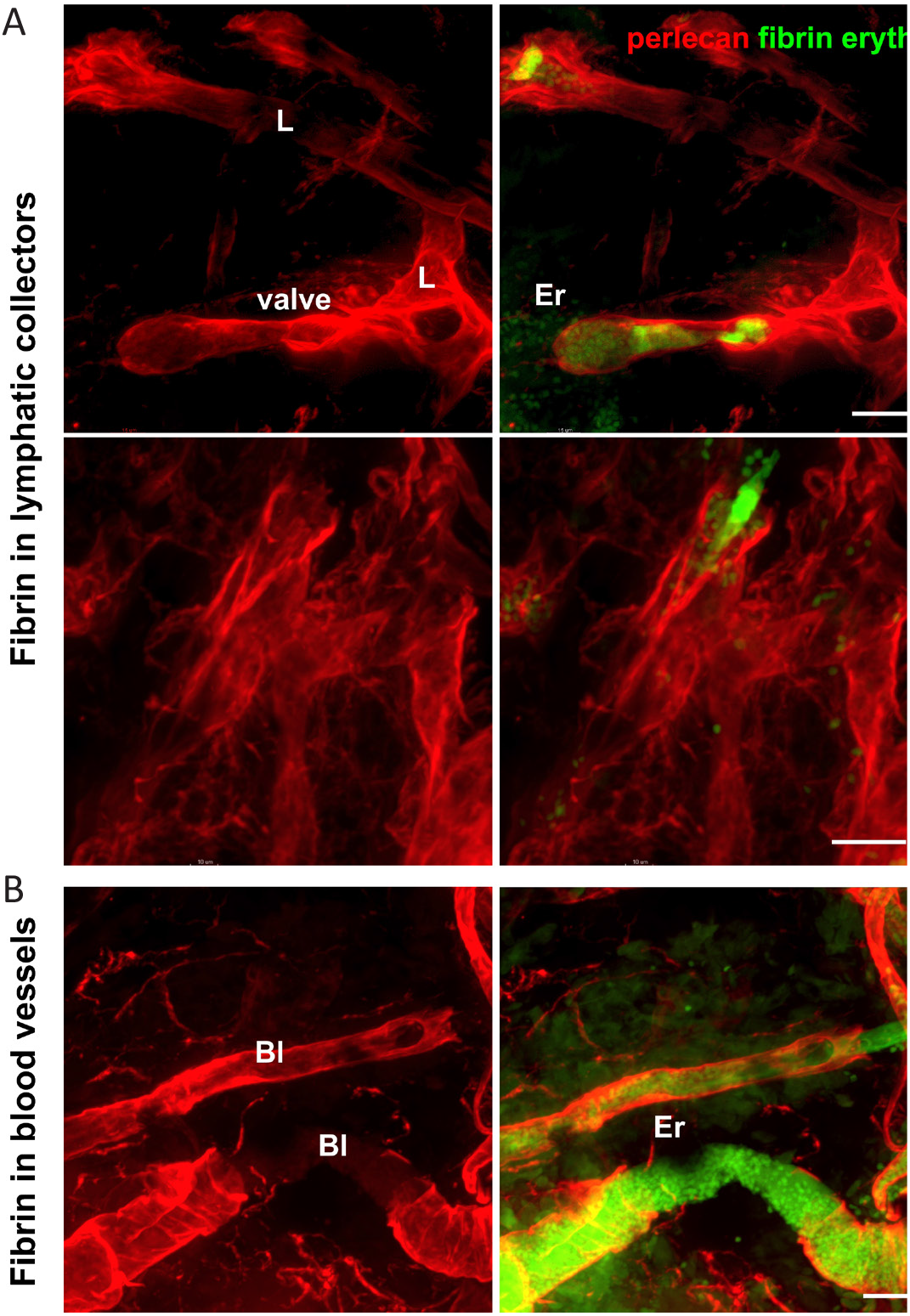
Fibrosis in peripheral lymphatics of B16/ F10 tumor implanted under the ear skin. A. Two examples of intralymphatic fibrosis identified at the edge of the lymphatic tumor. B16/F10 tumor cells were implanted for 7 days under the dorsal dermis of the ear. Green clots and autofluorescent lymphaticsArrows point to clusters of collagen IV-positive matrix inclusions inside lymphatic collecting vessels. Arrowheads point to disorganized lymphatic endothelial cells with absent cell-cell junctions. The intensity of collagen IV was normalized to the signal leveBl-blood vessels, Er-erythocytes, L-lymphatic collector. Scale bar: 50μm.

**Supplementary Fig. 2.**
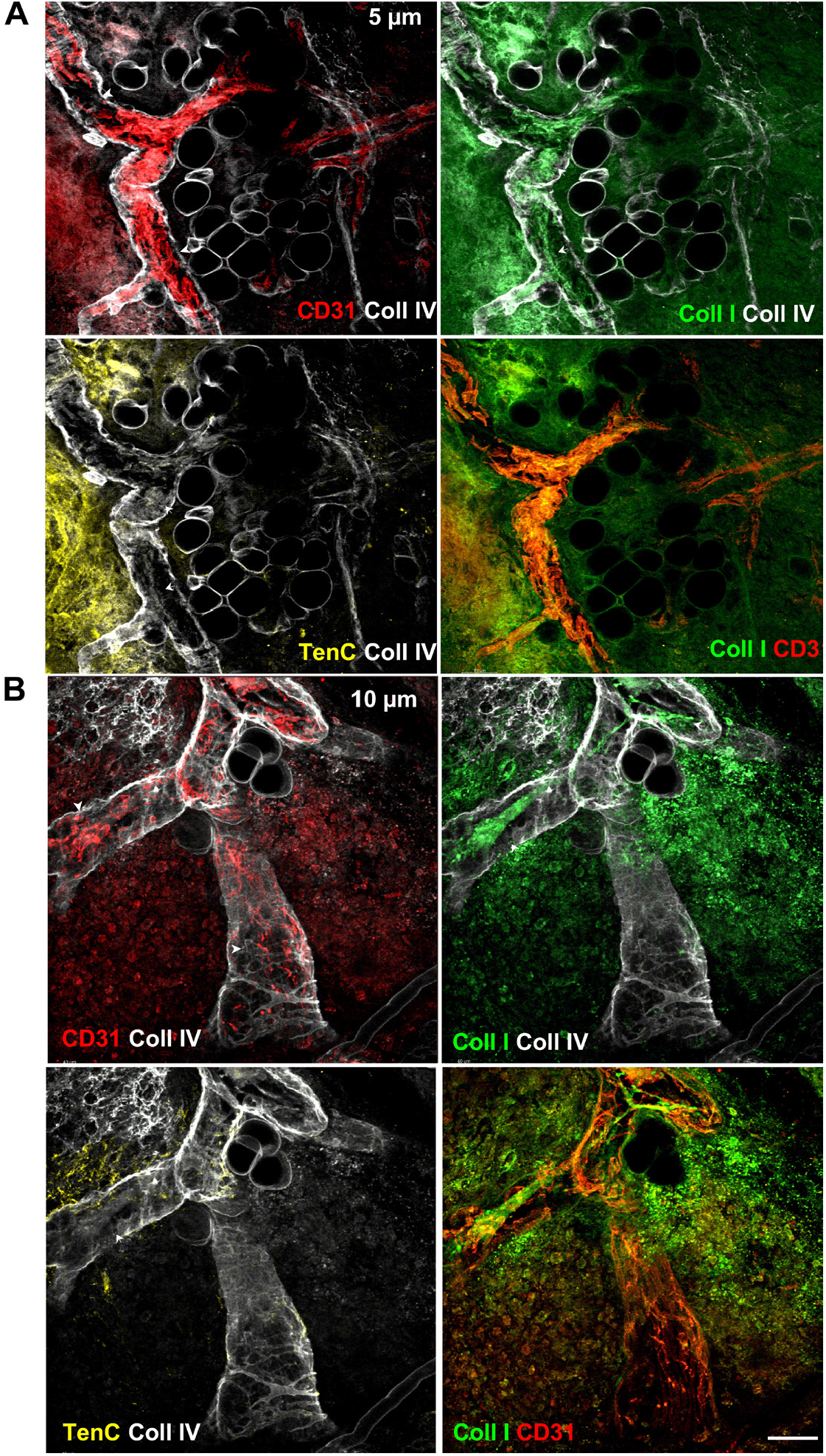
Lymph clots are forming within collectors at the periphery of B16/F10 melanoma. Three weeks after implantation of tumor melanoma cells tumor mass formed palpable solid mass under the dorsal dermis of a mouse ear. The mouse was i.v. injected with FITC-labelled fibrinogen and six hours later blood circulation was flushed with Ringer’s solution to remove blood and non-clottet fibrinogen from functional vessels. After that, the circulation was perfused with zinc-fixative, and the whole skin and tumor tissue were stained for basement membrane marker perlecan. Lymphatic collector vessels were identified by the presence of characteristic valves along which vessels change their diameter, from the narrow at the afferent tubular to wide at the efferent sinusoidal sides. Peritumoral lymphatic collectors had incomplete basement membrane and were filled with autofluorescent erythrocytes that were also present outside vessels in the tumor interstitium (top two rows). However, in contrast to non-perfused (non-functional) blood vessels (bottom row), erythrocyte presence did not overlap with clotted labeled fibrin. Scale bar: 25μm.

**Supplementary Fig. 3.**
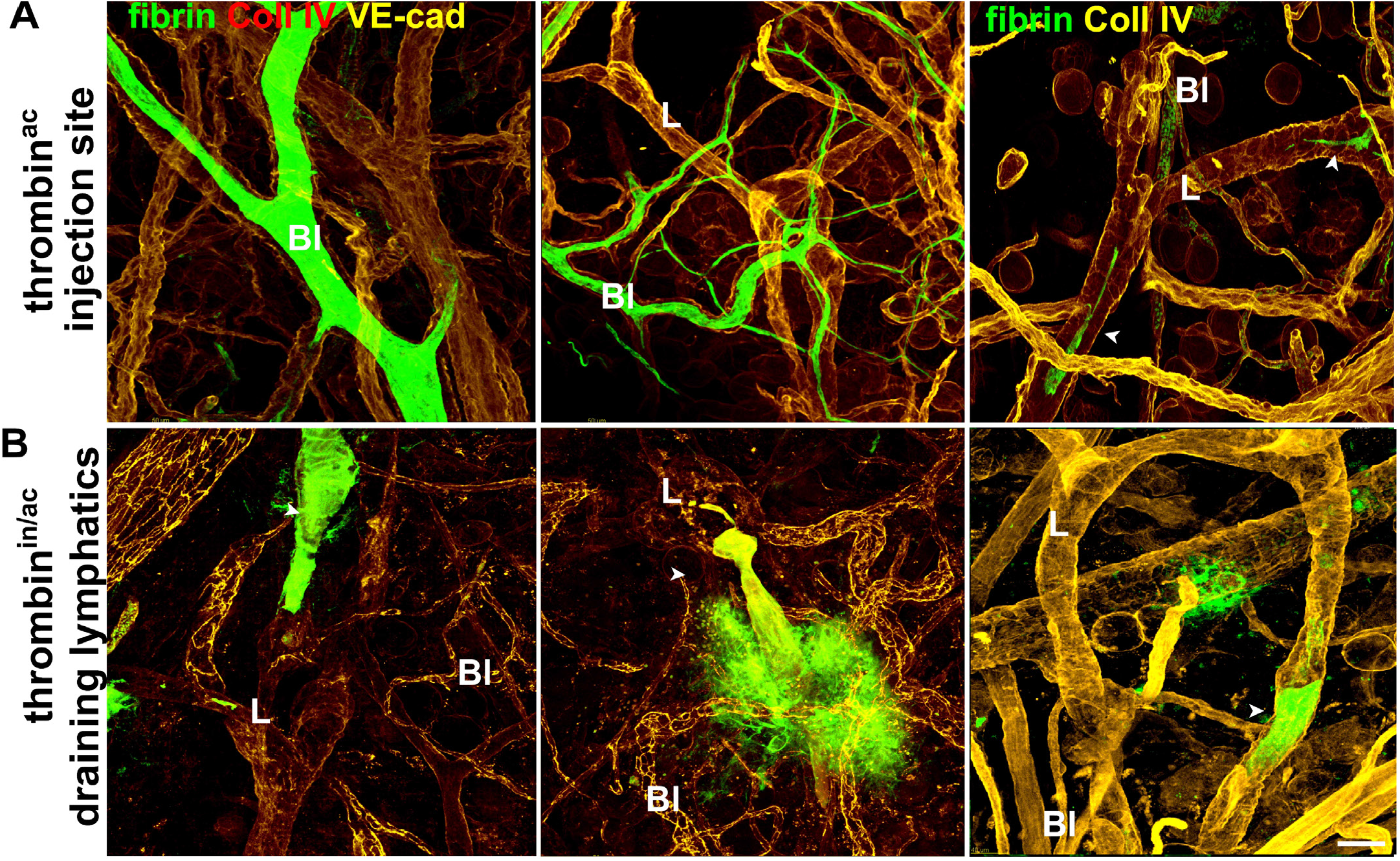
Comparison between active and partially active thrombin administration on intralymphatic and vascular blood clotting. Confocal microscopy of whole-mount ear. A. Site of Intradermal injection: Injection of concentrated active thrombin (thrombin^ac^) results in an instant but transient blood coagulation within blood vessels around the site of injection. Except for rare events, intralymphatic clotting was restricted only to the site around the injection area, and remote collectors were never occluded. Bl-blood vessels, L-lymphatics. B. Draining lymphatics: In contrast to active thrombin, two sequential injections of the mix of partially inactivated thrombin (1:6 active and inactive ratio, thrombin^in/ac^) had no effect on blood vessel coagulation and induced intralymphatic clotting only in remote collectors with no effect on blood coagulation at the site of injection. Intralymphatic clotting was often associated with lymphatic leakiness around the clot location (interstitial fibrin deposits in the bottom, middle image). Bl-blood vessels, L-lymphatic collectors. Scale bar: 50μm.

**Supplementary Fig. 4.**
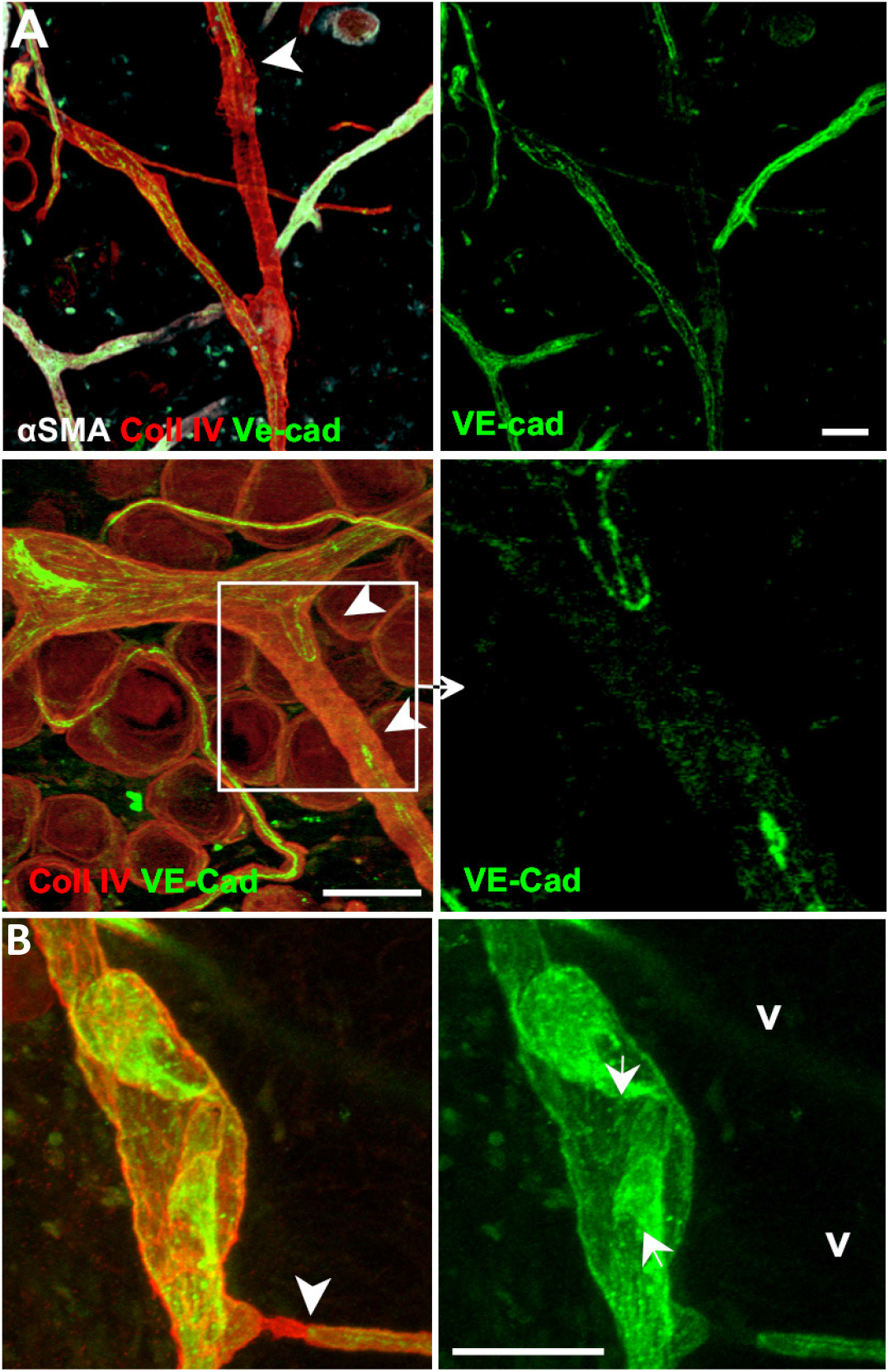
Epimorphic regeneration restored the function of decellularized lymphatics. **A**. Lymphatics repopulated the basement membranes of the original lymphatic collectors and did not sprout outside the lymphatic vasculature. Nine days after decellularization. Lymphatic endothelium sprouted along the basement membrane from opposite ends of surviving collectors completed repopulation of ghost vessels. During this regeneration phase, the ‘kissing’ connections (arrowheads) between advancing endothelial frontiers could be found within lymphatic ghosts. **B**. Even though lymphatic regeneration does not follow the random pattern of granulation growth, lymphatic endothelium re-growing along basement membranes of decellularized ghost vessels could still form abnormal structures, such as opposite-facing valves (arrows). Scale bar: 50μm.

**Supplementary Fig. 5.**
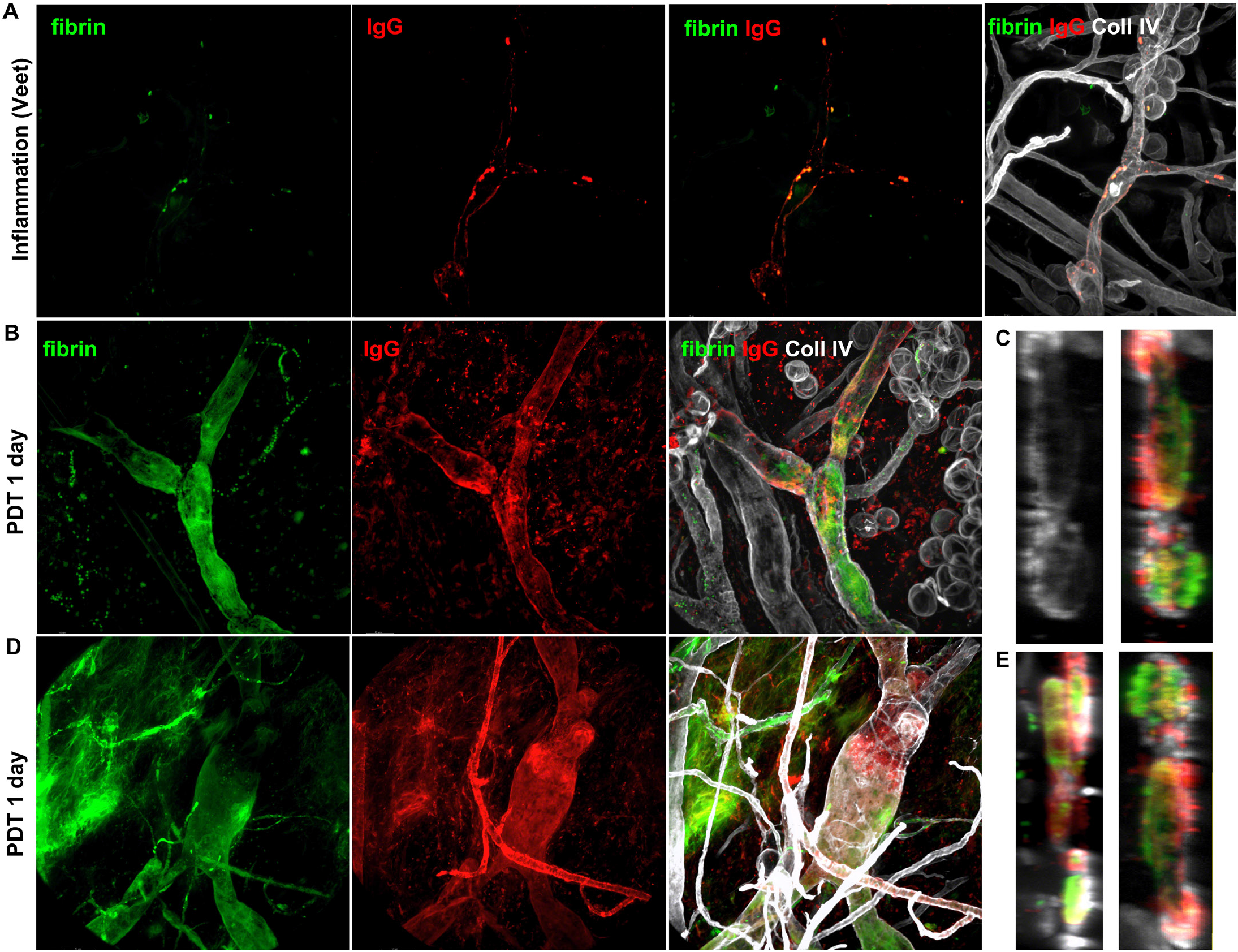
Dendritic cells populate regenerating ‘ghost vessels’ after lymphatic-specific photodynamic therapy. A. Epidermal Langerhans cells, classic antigen presenting cells in the ear. In the epidermis, Langerhans cells create a dense, 2-dimensional and dendrite interconnected network that stains strongly for MHCII and CD11c. Basement membrane (BM) is stained with collagen IV. B. In the control region of the ear treated with photodynamic therapy (PDT), lymphatics are not populated with antigen presenting cells. C. In contrast, lymphatic collectors decellularized with PDT are densely populated with mostly double-positive MHCII+ and CD11c+ antigen presenting cells even 10 days after the treatment. Hf-hair follicles. Scale bar: 50μm.

**Supplementary Fig. 6.**
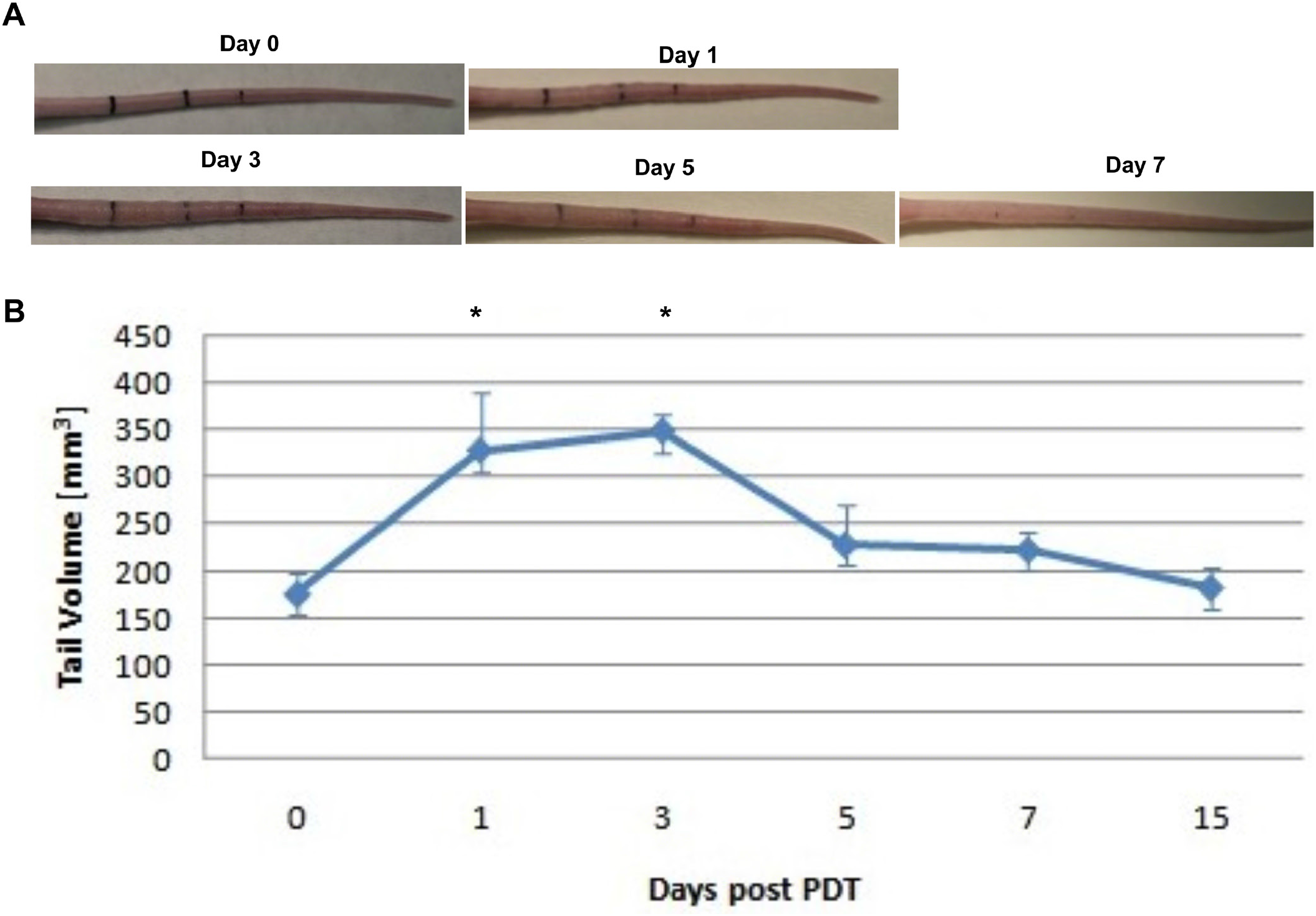
Lymphatic occlusion alone does not lead to persistent edema. Photodynamic therapy (PDT)-induced edema is transient and is resolved before lymphatic regenerate. 2 μg of Visudyne® (6x more than used to occlude the ear lymphatics) was injected in two sides of the tail with FITC-dextran. Draining lymphatics at a distance of 4 cm of the tail were irradiated with 25 J/cm^2,^ and they were imaged with IVIS every day for 15 days. A. Example tail before edema development (day 0), during its peak (day 1 to 3 after PDT) and after the edema resolution (day 5, 7). B. Graph representing the edema development as measured by tail volume, and resolution after PDT.

**Supplementary Fig. 7.**
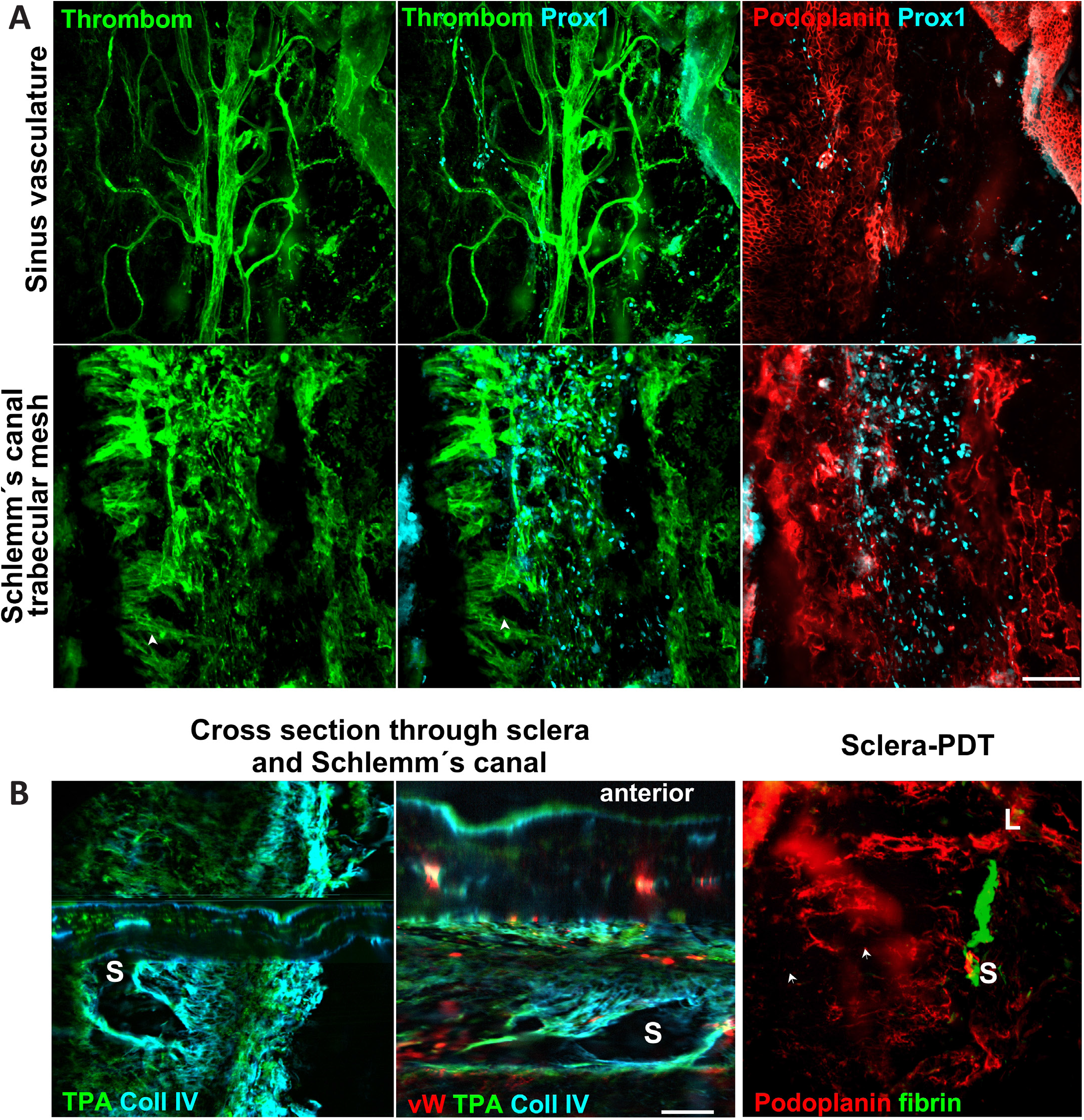
Decelularization of thrombomodulin-expressing endothelium of Schlemm canal leads to its thrombotic occultion. **A**. The location at the sclera-cornea junction, adjacent to the anterior chamber. Endothelium of trabecular meshwork (arrows) and Schlemm’s canal (S) stains for Prox1, a transcription factor expressed by lymphatics. Prox1 was not present in the cells lining trabecular meshwork, while podoplanin, a surface marker of skin lymphatics, stained trabecular meshwork, but not Schlemm’s canal. In contrast, endothelium lining Schlemm’s canal and its trabecular meshwork express thrombomodulin. B. Together with thrombomodulin, tissue plasminogen activator (TPA) expressed by the endothelium of Schlemm’s canal and trabecular network protect the canal from thrombotic occlusion. Cross-section through longitudinal plane of Schlemm’s canal depicts its lumen (S) and the lumen-occluding fibrin clot that formed after the destruction of the endothelium of Schlemm canal. Scale bar: 50μm.

## Notes

### Competing Interest Statement

The authors have declared no competing interest.

### Summary of Updates

Two authors, M. Swartz and A. Muchowicz ask to be removed from the Author list. Also, there are some minor corrections in the manuscript and Figures 1 and 3.

## References

1. Gimbrone (1987). “Vascular endothelium: Nature’s blood-compatible container.” Ann N Y Acad Scin

2. Rusznyák,& al. (1967). Lymphatics and lymph circulation. Physiology and pathology. Oxford, Pergamon Press.

3. Drinker,& al. (1933). Lymphatics, lymph and tissue fluid, The Williams & Wilkins Co.

4. Miller,& al. (2000). “Haemostatic factors in human peripheral afferent lymph.” Thromb Haemost

5. Opie (1913). “Thrombosis and occlusion of lymphatics.” J Med Res

6. Copley,& al. (1942). “Bleeding time, lymph time, and clot resistance in men.” J Clin Invest

7. Fantl,& al. (1953). “Coagulation in lymph.” J Physiol

8. Mayanskii,& al. (1966). “A comparative study of the clotting power of the blood and lymph.” Bulletin of Experimental Biology and Medicine

9. Le,& al. (1998). “Hemostatic factors in rabbit limb lymph: Relationship to mechanisms regulating extravascular coagulation.” Am J Physiol

10. Leak,& al. (2004). “Proteomic analysis of lymph.” Proteomics

11. Lippi,& al. (2012). “Hemostatic properties of the lymph: Relationships with occlusion and thrombosis.” Semin Thromb Hemost

12. Brinkman,& al. (1996). Vessel wall-mediated activation of the blood coagulation system. Vascular control hemost. VWM van Hinsberg. Amsterdam, the Netherlands, Harwood Academic Publishers: 85–106.

13. Drinker,& al. (1933). Inflammation. Lymphatics, lymph and tissue fluid, The Williams & Wilkins Co.

14. Aukland,& al. (1993). “Interstitial-lymphatic mechanisms in the control of extracellular fluid volume.” Physiol Rev

15. Wiig,& al. (2012). “Interstitial fluid and lymph formation and transport: Physiological regulation and roles in inflammation and cancer.” Physiol Rev

16. Engelmann,& al. (2013). “Thrombosis as an intravascular effector of innate immunity.” Nat Rev Immunol

17. Conway (2012). “Thrombomodulin and its role in inflammation.” Semin Immunopathol

18. Maruyama,& al. (1985). “Thrombomodulin is found on endothelium of arteries, veins, capillaries, and lymphatics, and on syncytiotrophoblast of human placenta.” J Cell Biol

19. Gould,& al. (1979). “The spandrels of san marco and the panglossian paradigm: A critique of the adaptationist programme.” Proc R Soc Lond B Biol Sci

20. Jafree,& al. (2021). “Mechanisms and cell lineages in lymphatic vascular development.” Angiogenesis

21. Kirchhofer,& al. (1994). “Endothelial cells stimulated with tumor necrosis factor-alpha express varying amounts of tissue factor resulting in inhomogenous fibrin deposition in a native blood flow system. Effects of thrombin inhibitors.” J Clin Invest

22. Laschinger,& al. (1990). “Production of plasminogen activator and plasminogen activator inhibitor by bovine lymphatic endothelial cells: Modulation by tnf-alpha.” Thromb Res

23. Drinker,& al. (1933). Lymph edema and elephantiasis. Lymphatics, lymph and tissue fluid, The Williams & Wilkins Co.

24. Olszewski (2002). “De novo lymph node formation in chronic inflammation of the human leg.” Ann N Y Acad Sci

25. Case,& al. (1991). “Vascular abnormalities in experimental and human lymphatic filariasis.” Lymphology

26. Fader,& al. (1986). “Evolution of lymph thrombi in experimental brugia malayi infections: A scanning electron microscopic study.” Lymphology

27. Hara,& al. (2013). “Lymphoedema caused by idiopathic lymphatic thrombus.” J Plast Reconstr Aesthet Surg

28. Cluzan (2007). “Peripheral lymphoedema treatment.” Phlebologie

29. Forster,& al. (2006). “Tissue factor and tumor: Clinical and laboratory aspects.” Clin Chim Acta

30. Ruf,& al. (2011). “Tissue factor and cell signalling in cancer progression and thrombosis.” J Thromb Haemost

31. Padera,& al. (2002). “Lymphatic metastasis in the absence of functional intratumor lymphatics.” Science

32. Idell (2003). “Coagulation, fibrinolysis, and fibrin deposition in acute lung injury.” Crit Care Med

33. Margaritescu (2015). Cutaneous metastases of breast carcinoma. Rare malignant skin tumors. F Rongioletti, I Margaritescu& al. New York, Springer: 313–316.

34. van der Meijden,& al. (2009). “Dual role of collagen in factor xii-dependent thrombus formation.” Blood

35. White-Adams,& al. (2010). “Laminin promotes coagulation and thrombus formation in a factor xii-dependent manner.” J Thromb Haemost

36. Shoskes (2011). Kidney transplant recipient surgery. New York, Springer.

37. Fahmy,& al. (2021). “Is microthrombosis the main pathology in coronavirus disease 2019 severity?-a systematic review of the postmortem pathologic findings.” Crit Care Explor

38. Bevilacqua,& al. (1984). “Interleukin 1 (il-1) induces biosynthesis and cell surface expression of procoagulant activity in human vascular endothelial cells.” J Exp Med

39. Lupu,& al. (2005). “Tissue factor-dependent coagulation is preferentially up-regulated within arterial branching areas in a baboon model of escherichia coli sepsis.” Am J Pathol

40. Liu,& al. (2004). “Thrombin and tumor necrosis factor alpha synergistically stimulate tissue factor expression in human endothelial cells: Regulation through c-fos and c-jun.” J Biol Chem

41. Kilarski,& al. (2013). “Intravital immunofluorescence for visualizing the microcirculatory and immune microenvironments in the mouse ear dermis.” PLoS One

42. van Hinsbergh (2012). “Endothelium--role in regulation of coagulation and inflammation.” Semin Immunopathol

43. Jesty,& al. (2005). “Positive feedbacks of coagulation: Their role in threshold regulation.” Arterioscler Thromb Vasc Biol

44. Mackman (2009). “The role of tissue factor and factor viia in hemostasis.” Anesth Analg

45. Esmon (1995). “Thrombomodulin as a model of molecular mechanisms that modulate protease specificity and function at the vessel surface.” FASEB J

46. Fuentes-Prior,& al. (2000). “Structural basis for the anticoagulant activity of the thrombin-thrombomodulin complex.” Nature

47. Laura,& al. (1980). “(p-amidinophenyl)methanesulfonyl fluoride, an irreversible inhibitor of serine proteases.” Biochemistry

48. (2019). Hemostasis and thrombosis. Cham, Switzerland, Springer Nature Switzerland.

49. Chen,& al. (2010). “Sterile inflammation: Sensing and reacting to damage.” Nat Rev Immunol

50. Esmon (2010). “Far from the heart: Counteracting coagulation.” Nat Med

51. Wu,& al. (1996). “Role of endothelium in thrombosis and hemostasis.” Annu Rev Med

52. Egorina,& al. (2008). “Granulocytes do not express but acquire monocyte-derived tissue factor in whole blood: Evidence for a direct transfer.” Blood

53. Wang,& al. (2005). “Effects of factor ix or factor xi deficiency on ferric chloride-induced carotid artery occlusion in mice.” J Thromb Haemost

54. Kilarski,& al. (2014). “Optimization and regeneration kinetics of lymphatic-specific photodynamic therapy in the mouse dermis.” Angiogenesis

55. Inai,& al. (2004). “Inhibition of vascular endothelial growth factor (vegf) signaling in cancer causes loss of endothelial fenestrations, regression of tumor vessels, and appearance of basement membrane ghosts.” Am J Pathol

56. van Meeteren,& al. (2012). “Regulation of endothelial cell plasticity by tgf-beta.” Cell Tissue Res

57. Guc,& al. (2014). “Long-term intravital immunofluorescence imaging of tissue matrix components with epifluorescence and two-photon microscopy.” J Vis Exp

58. Randolph,& al. (2005). “Dendritic-cell trafficking to lymph nodes through lymphatic vessels.” Nat Rev Immunol

59. Drinker,& al. (1933). Relation of the lymphatics to edema and the consequences of lymphatic obstruction. Lymphatics, lymph and tissue fluid, The Williams & Wilkins Co.

60. Benichou,& al. (2011). “Immune recognition and rejection of allogeneic skin grafts.” Immunotherapy

61. Lund,& al. (2016). “Lymphatic vessels, inflammation, and immunity in skin cancer.” Cancer Discov

62. Lakkis,& al. (2000). “Immunologic ‘ignorance’ of vascularized organ transplants in the absence of secondary lymphoid tissue.” Nat Med

63. Yamagami,& al. (2001). “The critical role of lymph nodes in corneal alloimmunization and graft rejection.” Invest Ophthalmol Vis Sci

64. Fulmer,& al. (1963). “Transplantation of cardiac tissue into the mouse ear.” Am J Anat

65. McFarland,& al. (2009). “Skin allograft rejection.” Curr Protoc Immunol

66. Ruddle (2014). “Lymphatic vessels and tertiary lymphoid organs.” J Clin Invest

67. Gorecki,& al. (1991). “Evidence that liposome incorporation of cyclosporine reduces its toxicity and potentiates its ability to prolong survival of cardiac allografts in mice.” Transplantation

68. Colvin,& al. (2005). “Antibody-mediated organ-allograft rejection.” Nat Rev Immunol

69. Menkin (1931). “Studies on inflammation : Vii. Fixation of bacteria and of particulate matter at the site of inflammation.” J Exp Med

70. Menkin (1931). “Studies on inflammation : Vi. Fixation of trypan blue in inflamed areas of frogs.” J Exp Med

71. Menkin (1931). “Studies on inflammation : V. The mechanism of fixation by the inflammatory reaction.” J Exp Med

72. Guytron,& al. (2016). The body fluids and kidneys. Textbook of medical physiology. Philadelphia, Elsevier.

73. Kilarski (2018). “Physiological perspective on therapies of lymphatic vessels.” Adv Wound Care (New Rochelle)

74. Medawar (1948). “Immunity to homologous grafted skin; the fate of skin homografts transplanted to the brain, to subcutaneous tissue, and to the anterior chamber of the eye.” Br J Exp Pathol

75. Simpson (2006). “A historical perspective on immunological privilege.” Immunol Rev

76. Aspelund,& al. (2015). “A dural lymphatic vascular system that drains brain interstitial fluid and macromolecules.” J Exp Med

77. Barker,& al. (1968). “The role of afferent lymphatics in the rejection of skin homografts.” J Exp Med

78. Lambert,& al. (1965). “The role of the lymph trunks in the response to allogeneic skin transplants.” Transplantation

79. Joffre,& al. (2008). “Prevention of acute and chronic allograft rejection with cd4+cd25+foxp3+ regulatory t lymphocytes.” Nat Med

80. Lorentz,& al. (2011). “Engineered aprotinin for improved stability of fibrin biomaterials.” Biomaterials

81. Pepper,& al. (2015). “A prevascularized subcutaneous device-less site for islet and cellular transplantation.” Nat Biotechnol

82. Ferri (2014). Ferri’s clinical advisor 2013: 5 books in 1. Philadelphia, PA, Elsevier Health Sciences.

83. Menkin (1960). “Biochemical mechanisms in inflammation.” Br Med J

84. Rutkowski,& al. (2006). “Characterization of lymphangiogenesis in a model of adult skin regeneration.” Am J Physiol Heart Circ Physiol

85. Rockson (2021). “Lymphedema, inflammation, and fat.” Lymphat Res Biol

86. Chakraborty,& al. (2013). “Lymphatic filariasis: Perspectives on lymphatic remodeling and contractile dysfunction in filarial disease pathogenesis.” Microcirculation

87. Patel,& al. (2015). “Lymphatic mapping and lymphedema surgery in the breast cancer patient.” Gland Surg

88. Olszewski,& al. (2015). “A novel method of edema fluid drainage in obstructive lymphedema of limbs by implantation of hydrophobic silicone tubes.” J Vasc Surg Venous Lymphat Disord

89. Thomson,& al. (2014). “A lymphatic defect causes ocular hypertension and glaucoma in mice.” J Clin Invest

90. Palumbo,& al. (2002). “Spontaneous hematogenous and lymphatic metastasis, but not primary tumor growth or angiogenesis, is diminished in fibrinogen-deficient mice.” Cancer Res

91. Tammela,& al. (2011). “Photodynamic ablation of lymphatic vessels and intralymphatic cancer cells prevents metastasis.” Sci Transl Med

92. Loo,& al. (2021). “Covid-19, immunothrombosis and venous thromboembolism: Biological mechanisms.” Thorax

93. Achar,& al. (2020). “Covid-19-associated neurological disorders: The potential route of cns invasion and blood-brain relevance.” Cells

